# Synthesis of substituted oxabicyclic phenol derivatives with increased affinity and activity for the classical estrogen receptor ERα

**DOI:** 10.64898/2026.09.25.754525

**Authors:** Richard A. Pepermans, Antonio J. Tenorio, Travis Saxon, Marat R. Talipov, William A. Maio, Chinnasamy Ramesh, Eric R. Prossnitz, Jeffrey B. Arterburn

## Abstract

The classical estrogen receptor α (ERα) is a steroid hormone receptor known for its role in both health and disease. ERα, like its counterpart ERβ, acts as a receptor for its endogenous ligand 17β-estradiol (E2) and is capable of binding a diverse set of natural and synthetic small molecules. Several screening campaigns have identified various ER ligands, based on the oxabicyclo[3.3.1]-nonene scaffold, that act as ER agonists. One such campaign extensively characterized an oxabicyclic ER agonist, termed AB-1. Herein, we report the synthesis of a series of analogs based on the scaffold of AB-1 (herein termed **1**), focusing on the enhancement of their binding to ERα and functional potencies in ER-mediated assays. We identify a new substituted compound **9** that achieves subnanomolar potencies for ERα binding (IC_50_= 0.45 nM) and ER-dependent transcription (EC_50_= 0.37 nM) and proliferation (EC_50_= 0.03 nM), with potencies approaching those of **E2**. The high potency of **9** provides a promising scaffold for the potential development of new ERα-targeted therapeutics.

## 1. Introduction

The classical estrogen receptors α (ERα) and β (ERαβ) are ligand-binding nuclear hormone receptors that play a role in various physiological settings. High structural homology, particularly in the ligand-binding cavity between ERα and ERβ, partially overlapping tissue expression and similar affinities for their natural ligand, 17β-estradiol (E2), made dissecting their individual roles challenging [1–4]. Nevertheless, compounds were discovered that display preferential binding to the individual subtypes. Such compounds include the natural products isoflavone, genistein and other phytoestrogens, as well as synthetic compounds, such as PPT and DPN, [2, 5–7].

In the context of breast cancer, which is often driven by aberrant ERα signaling, ERα has been a therapeutic target for decades. Drugs such as selective estrogen receptor modulators/degraders (SERMs/SERDs) have revolutionized patient outcomes, prolonging the lives of millions of women [8–11]. Unfortunately, many women eventually relapse of their disease, with tumors typically becoming resistant to the original therapy [12–14]. This emboldens the ongoing search for new high affinity ER ligands that could potentially be converted into future ERα-targeting therapeutics such as SERDs or the more recently explored ERα proteolytic-targeting chimeras (PROTACs) [15–17].

ER ligands display various structures, typically harboring planar structures containing several hydrocarbon rings similar to that of E2. Of interest, several independent screening campaigns identified an oxabicyclic molecular scaffold, particularly the oxabicyclo[3.3.1]-nonene scaffold, as a high affinity ER-binding template, with compounds harboring this scaffold generally acting as potent ER agonists [18–20]. While these screening studies reported minimal characterization of the compounds, they observed moderate to high ERβ selectivity (over ERα) of the compounds. In a separate screening project, we identified an oxabicyclo[3.3.1]-nonene compound, termed AB-1, that was also identified in other independent screens, as an ER agonist [21]. However, contrary to previous findings, we observed low to moderate ERβ selectivity of AB-1 in our characterization of the compound.

In an attempt to identify new high affinity ER ligands that could potentially serve as the basis for future ERα therapeutics, we set out to identify novel, high affinity ERα ligands through structure-activity relationship studies. Since we previously characterized the oxabicyclic ERα agonist AB-1 [21], we selected it (herein termed **1**) as the starting point for structural modifications with the goal of identifying molecules with subnanomolar ERα-binding affinity. The current investigation was motivated by recognizing the need to both refine and optimize structure-activity features associated with ER-binding affinities and potencies, and to provide detailed experimental procedures and yields for optimized compounds to facilitate further studies, since these important details were largely unavailable for compounds described in previous screening studies [18–20]. Our synthetic rationale for introducing structural modifications was driven by recognition of the important molecular features identified in **Figure 1a**. The hydrogen-bonding phenol (**A**) and primary alcohol (**B**) functional groups occur in the endogenous ligand **E2**, as well as many other estrogenic compounds, and structural studies have demonstrated that these form strong interactions with protein residues in the ER ligand binding site [7, 22]. The introduction of fluorine in the ortho-position of a phenol group provides a way to increase the acidity and alter H-bonding interactions through electronic effects without significantly increasing steric volume. A multicomponent approach to generate dienophiles from 1,3-dicarbonyl precursors provides an efficient pathway for introducing other functional groups with altered steric and electronic properties in region **B**. We also envisioned modification of the conformational and steric properties of the unsaturated bicyclic ether scaffold in region **C** through introduction of methyl substituents and hydrogenation of the cyclohexenyl group. Herein, we identify a lead compound **9** possessing a novel saturated oxabicyclo[3.3.1]nonan-2-yl scaffold that exhibits a 90-fold improvement in ERα binding affinity compared to the parent compound AB-1 (**1**).

**Fig 1.**
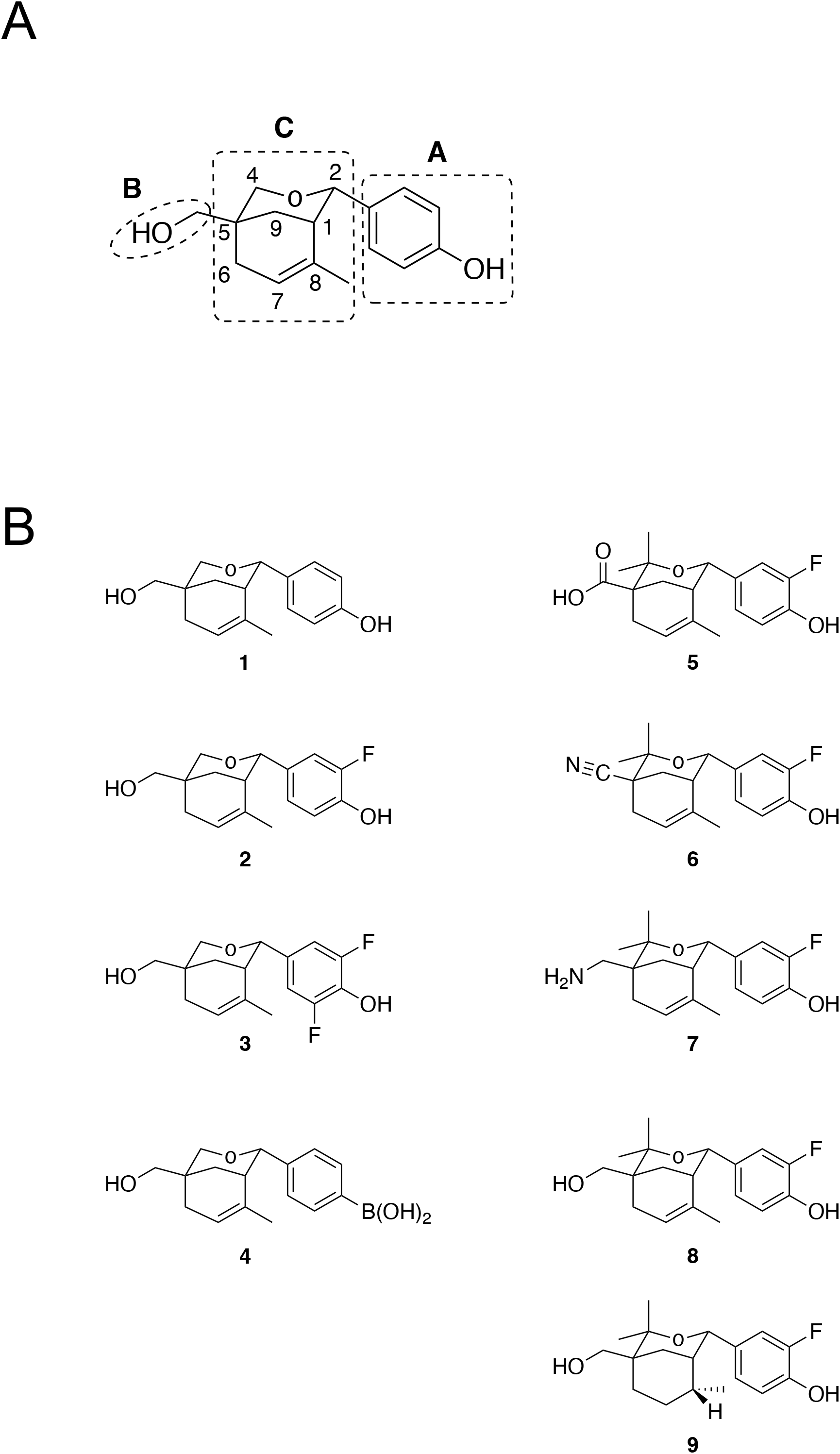
(A) Regions of interest for structure-activity exploration of parent compound **1**. (B) Structures of analog series derived from **1**.

## 2. Results and Discussion

### 2.1. Compound series design and synthesis

Based on our initial study of **1** [21] and informed by other results from independent screening studies that had identified related oxabicyclo[3.3.1]nonene derivatives as ERα/β agonists [19], we set out to design and synthesize a focused series of analogs to enable comparison of their ERα/β binding and receptor-mediated activities. Previous investigations had focused on identifying structural features of substituted derivatives associated with general increased ER subtype selectivity [18–20]. Our interests were directed towards optimizing the structure of analogs within the oxabicyclo[3.3.1]nonene scaffold to increase binding affinity specifically for ERα, relative to **1**. We also recognized the need for providing efficient, scalable synthetic procedures and spectroscopic characterization of these analogs, since full experimental details and characterization data are not available for most related published compounds [18–20].

Our previously reported synthesis of **1** used a multicomponent reaction in which catalytic zinc(II) chloride was used to initiate the condensation reaction between the 1,3-dicarbonyl active methylene compound diethylmalonate and formaldehyde to generate the dienophile intermediate, diethylmethylene malonate, which subsequently undergoes a Zn(II)-catalyzed Diels-Alder cyclization with the isoprene diene [21]. This *in situ* approach for generating the dienophile was advantageous by avoiding the need to use highly electrophilic and polymerization-sensitive reagents, which have limited commercial availability. The inclusion of other functionalized 1,3-activated methylene compounds in this approach was expected to enable access to other substituted products with varying H-bond donor and acceptor groups. We had also evaluated a variety of Lewis acid catalysts for the critical Prins cyclization reaction that incorporated 4-hydroxybenzaldehyde to form the oxabicyclic scaffold, and identified hafnium(IV) triflate as the most effective catalyst. Our plan for initial structure-activity studies identified possible modifications in regions A–C of parent compound **1** (**Fig. 1a**).

Following our general synthetic route, substitutions in the phenolic group in region A at the 2-position of the scaffold would be accessible by varying the benzaldehyde reagent in the Prins cyclization reaction step. The hydroxymethyl group at the 5-position in region B could potentially be altered through incorporation of different functional groups in the dienophile, as could the substituents on the 4-postion of region C adjacent to the oxygen in the pyranose ring. We designed the specific set of substituted analogs **2-9** for our initial screening studies (**Fig. 1b**).

The syntheses of compounds **2-4**, possessing substitutions in the phenolic region A, were accomplished starting from diethylmalonate as shown in **Schemes 1-2**. The Zn(II)-catalyzed condensation and Diels-Alder reaction produced the substituted cyclohexene **10** and subsequent reduction of the ester groups with lithium aluminum hydride provided the diol **11**. The 2-fluoro (**2**) and 2,6-difluoro (**3**) compounds were obtained directly through a Hf(IV)-catalyzed Prins cyclization reaction of diol **11** with 3-fluoro-4-hydroxybenzaldehyde or 3,5-difluoro-4-hydroxybenzaldehyde, respectively (**Scheme 1**).

The phenylboronic acid functional group in **4** was introduced as a potential isosteric analog of the phenol group [23]. We prepared **4** as shown in **Scheme 2**. Starting from the previously described diol **11**, the Prins cyclization reaction was carried out to incorporate a 4-formylboronic acid derivative that was protected as an *N*-methyliminodiacetic acid (MIDA) boronate **12**. The MIDA group masks the electrophilicity of the boronic acid group and facilitates solubility in the acetonitrile reaction media, enabling the Hf(IV)-catalyzed Prins cyclization reaction to provide **13**. The MIDA protecting group in **13** was easily removed by mild hydrolysis with sodium bicarbonate to afford the product boronic acid **4**. The syntheses of **5** and **8** proceeded from the multicomponent Diels-Alder reactions of the *tert*-butyl and ethyl esters of acetoacetic acid, respectively (**Scheme 3**). The multicomponent Diels-Alder procedure, starting from the ethyl and *tert*-butyl acetoacetate esters, gave the corresponding cyclohexene products **14a** and **14b**. The methyl ketone group underwent selective addition of the nucleophilic Grignard reagent methylmagnesium iodide to provide the tertiary alcohol products **15a** and **15b**, while leaving the ester groups intact. The Prins cyclization reaction of ethyl ester **15a** gave the desired bicyclic product **16a**. However, multiple attempts to hydrolyze the ethyl ester group in **16a** to give the carboxylic acid derivative **5** were unsuccessful, even under the forcing conditions of high temperatures and strong base or acid. The lack of hydrolytic reactivity of the ethyl ester in **16a** is attributed to the unfavorable steric interactions between the tetrahedral intermediate and the adjacent bicyclic scaffold that would occur during carbonyl hydrolysis, increasing the energy required for this normally facile process. The desired compound **8**, possessing a primary alcohol at position 5, was obtained by reduction of the ethyl ester in **16a** with lithium aluminum hydride. An alternative route was developed for production of the carboxylic acid derivative **5** starting from *tert-*butyl acetoacetate and proceeding through the analogous process of Zn(II)-catalyzed multicomponent condensation/Diels-Alder reaction to produce the cyclohexene **14b**, followed by selective addition of the methylmagnesium iodide Grignard reagent to provide the tertiary alcohol **15b**. The acid-sensitive *tert*-butyl ester group in **15b** was removed by hydrolysis with trifluoroacetic acid in DCM to give the β-hydroxycarboxylic acid **15c**. The Hf(IV)-catalyzed Prins cyclization reaction of **15c** with 3-fluoro-4-hydroxybenzaldehyde then gave the desired carboxylic acid analog **5**.

The saturated analog **9** was obtained by selective hydrogenation of the cyclohexene ring in **8** using 10% Pd/C catalyst with H_2_ in ethanol as shown in **Scheme 3**. The hydrogenation was observed to be highly selective giving a single diastereomer. This finding revealed an interesting stereochemical feature that was observed through inspection of selected signals in the preceding ^1^H NMR spectra. The C8-methyl group in the cyclohexene intermediates **14a** and **15a** occur at δ 1.62 and δ 1.61 in CDCl_3_, respectively, which are typical for allylic methyl groups. However, following the Prins cyclization reaction, the C8-methyl group in **16a** was observed to have shifted significantly upfield, occurring at δ 0.95 in CDCl_3_. This upfield shift is attributed to the orientation of the C8-methyl group towards the planar face of the aromatic ring where anisotropic shielding effects can occur. The reduction of the ester group in **16a** with lithium aluminum hydride provides the alcohol **8** (**Scheme 3**), which exhibits the C8-methyl group ^1^H NMR signal at δ 0.91 in acetone-d_6_. Catalytic hydrogenation of **8** produces the saturated **9** as a single diastereomer, in which this C8-methyl group ^1^H NMR signal is observed even further upfield at δ 0.34 in acetone-d_6_, where it now appears as a doublet (*J* = 10.0 Hz) due to spin splitting with the adjacent cyclohexyl hydrogen atom. This increased shielding of the C8-methyl group in **9** is attributed to increased proximity within the anisotropic shielding region generated by the phenol. The diastereoselectivity of this hydrogenation can be rationalized by steric effects of the 4,4-dimethyl and phenol groups blocking access to the internal face of the alkene, leaving the external alkene face available for interaction with the hydrogenation catalyst.

The commercial availability of ethyl 2-cyanoacrylate, the active ingredient in many commercial instant glues, provides convenient access for using this compound directly as a dienophile in our synthetic approach as shown in **Scheme 4**. The Zn(II)-catalyzed Diels-Alder reaction of this reagent with isoprene was effective at ambient temperature and afforded the cyclohexene compound **17**. An excess (5 equiv.) of methylmagnesium iodide Grignard reagent provided selective conversion of the ethyl ester group to the desired tertiary alcohol derivative **18**, leaving the nitrile group intact. Compound **18** then underwent the Hf(IV)-catalyzed Prins cyclization reaction with 3-fluoro-4-hydroxybenzaldehyde to provide the desired bicyclic nitrile analog **6** (**Scheme 4**). The reduction of the nitrile group with lithium aluminum hydride gave **7**, which possesses a primary amine group at the 5-position.

### 2.2. Compound series assessment for ERα/β-LBD binding affinity

We initiated the assessment of synthetic compounds **1**-**9**, using a TR-FRET competitive binding assay employing a fluorescent estrogen derivative, to evaluate the structural effects of the various molecular substitutions on the binding affinities of the compounds to the ligand-binding domain of human ERα and ERβ. The binding affinities of the compounds, as well as E2 as a reference, are expressed as IC_50_ values and are given, along with their corresponding binding curves, in **Figure 2**.

**Fig 2.**
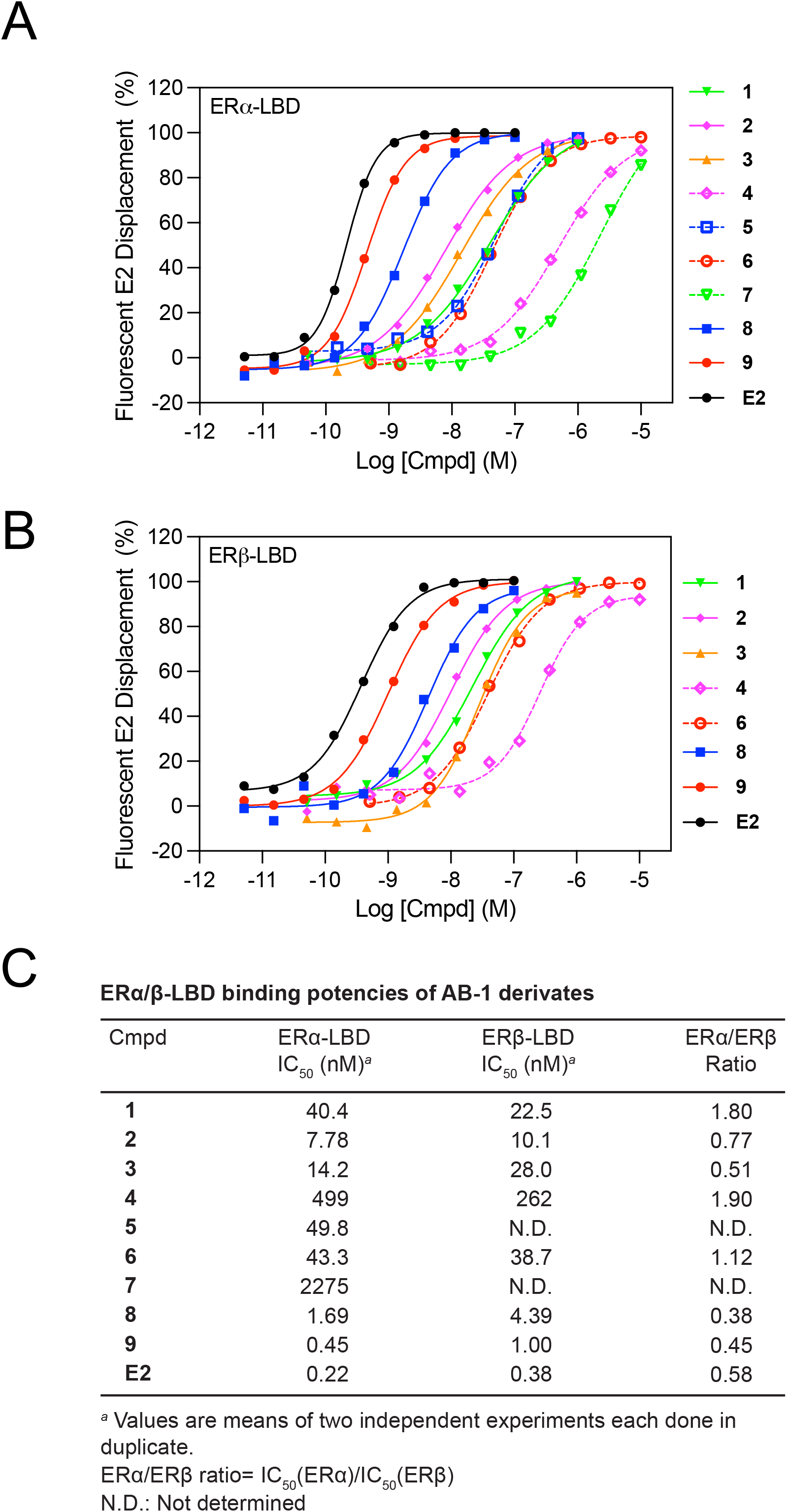
Binding affinities of compounds **1**-**9** and **E2** for ERα-LBD and ERβ-LBD. Dose-dependent displacement of a fluorescent E2 analogue from ERα-LBD (A) and ERβ-LBD (B) by the indicated compounds. (C) IC_50_ values of the dose-response curves shown in (A) and (B) and their corresponding ER-subtype binding ratio. Fluorescent E2 displacement is shown in (A) and (B) as a percentage compared to negative (1 % DMSO (v/v)) and positive (1 μM E2 in 1 % DMSO (v/v)) controls. Values are the means of two independent experiments each performed in duplicate.

Many synthetic ER ligands commonly share a phenolic ring that mimics the A-ring in E2, and other natural estrogens, whose phenol forms important H-bonds with the residues E353 and R394 and a water molecule in the ERα-LBD [22]. Furthermore, a crystal structure of **1** bound to the ERα-LBD in complex with a GRIP1 co-activator peptide reveals a similar H-bond network, exhibiting direct H-bond formation between the phenol in region A of **1** (**Fig. 1a**) and E353 of ERα [20]. These observations emphasize the importance of this phenol group in **1**, and we therefore first evaluated the effects of substitutions in region A of **1**, while preserving the phenol.

Addition of an *ortho*-fluoride group in **2**, resulted in a ∼5-fold increase in ERα binding affinity from IC_50_= 40.4 nM (**1**) to IC_50_=7.78 nM (**2**) (**Fig. 2a and c**). A modest ∼2-fold increase in ERβ binding from IC_50_= 22.5 nM to IC_50_= 10.1 nM was observed with the *ortho*-fluoride substituent. It is worth noting that **2** was previously identified in an independent screening campaign (therein termed compound 10) as an agonist of ERα, but its binding affinity for the receptor was not assessed [19]. The addition of two *ortho*-fluorides in **3** gave a ∼3-fold improvement in ERα binding compared to **1**, but a ∼2-fold decrease compared to **2**, with minimal effect on ERβ binding affinity compared to **1**. Recognizing the critical role of the phenol ring in **1** for efficient binding to ERα, we also evaluated replacement of the ring with a phenylboronic acid substituent in **4**. This group has been used as an isostere of the phenol group and found to improve oral bioavailability, while maintaining high ERα binding affinity in a boronic acid analog of the potent ERα antagonist fulvestrant [23]. Here, the introduction of the boronic acid substituent dramatically weakened ER binding capability, resulting in ∼12-fold reductions in both the ERα and ERβ binding affinities of **4**. A crystal structure of **1** bound to ERα shows a H-bond between the phenol of **1** and E353 in ERα [20]. The larger boronic acid group in **4** may impede binding and disrupt effective H-bond formation between the phenol in **4** and E353 in ERα, which would explain the loss in binding affinity. Based on the modification explored in region A of **1**, the single *ortho*-fluoride group exhibits the highest increase in ERα binding affinity and was maintained in the remaining analogs in our series.

We synthesized analogs **5-8** (**Fig. 1b**) by the routes shown in **Schemes 3 and 4**. A previous SAR study found that incorporating 4,4-dimethyl substituents improves the potency of **1** by 80-fold in an ER transcription assay and therefore this modification was included in the design of compounds **5**-**8** [19]. Crystal structure studies demonstrate that the hydroxymethyl group in region B of **1** enhances binding of the compound to ERα-LBD through formation of an H-bond with residue H524 in ERα [20]. This mimics the H-bond formed between the 17β-OH of E2 with ERα [22]. We sought to gain insight into the effects of other functional groups with different H-bond donor/acceptor properties through comparison of the binding affinities with compounds **5-8** using the data presented (**Figure 2**).

The analogs possessing a carboxylic acid **5** or nitrile group **6** (in addition to the *ortho*-fluoride and 4,4-dimethyl substitutions) failed to exhibit improvements in ERα binding affinities and gave IC_50_= 49.8 nM and IC_50_= 43.3 nM respectively, which are comparable to that of parent compound **1**. The ionized carboxylate group in **5** would experience increased solvation and is only capable of functioning as an H-bond acceptor, and therefore, these attributes failed to improve binding. No improvement in ERα or ERβ binding affinities were observed with the nitrile group in **6**. The linear, sp-hybridized nitrile group is neutral, but lacks H-bond binding properties. The amine analog **7** exhibited a large ∼56-fold decrease in ERα binding affinity compared to **1**. In the buffered binding assay conditions, the amine group in **7** would be ionized to the positively charged ammonium salt, which could function as a H-bond donor, but would also experience increased solvation. Compound **8** exhibited ∼24-fold increased ERα binding affinity and ∼5-fold improvement in ERβ binding relative to **1**, demonstrating the desirable features of the neutral, H-bond donating hydroxymethyl group within the relatively hydrophobic binding pocket in ERα-LBD. Of note, **8** was previously identified in an independent screening campaign (therein termed compound 42) as an agonist of ERα, but its binding affinity for the receptor was not assessed [19].

The oxabicyclic scaffold of **1** interacts weakly with L384 in ERα through van der Waals contacts [20]. We hypothesized that the changes in bond angles and molecular volume that would accompany saturation of the planar sp^2^ hybridized alkene group in region C of **1** would modify the weak hydrophobic interactions and affect the binding affinity. This modification was achieved by palladium-catalyzed hydrogenation of **8**. The resulting compound **9** exhibited subnanomolar binding affinity for ERα with IC_50_= 0.45 nM and near subnanomolar affinity for ERβ with an IC_50_= 1.00 nM. This represents increases in binding affinities for ERα and ERβ relative to the parent compound **1** of ∼90-fold and ∼22-fold, respectively, and a 4-fold improvement compared with **8**. Of note, the preferential binding to ERβ observed for parent compound **1** is reversed in compounds **8** and **9**, with both analogs now showing slight binding preference for ERα.

### 2.3. Compound effects on cell proliferation and viability

We next assessed how the molecular modifications to parent compound **1** affected the potency of the derivative compounds in functional cellular assays. For our initial functional assessment, we evaluated cell proliferation and viability of the E2-sensitive MCF-7 cell line, which upon ERα activation (with E2 or other ERα agonists) induces increased proliferation and viability, employing the CellTiter-Glo luminescence-based viability assay, which quantitates cellular ATP as a reflection of both cell proliferation and viability. The potencies and dose-response curves of **1**-**9** in the MCF-7 viability assay are given in **Figure 3a and b**. Due to their poor ERα binding affinities, **4** and **7** were not assessed in this or subsequent assays.

**Fig 3.**
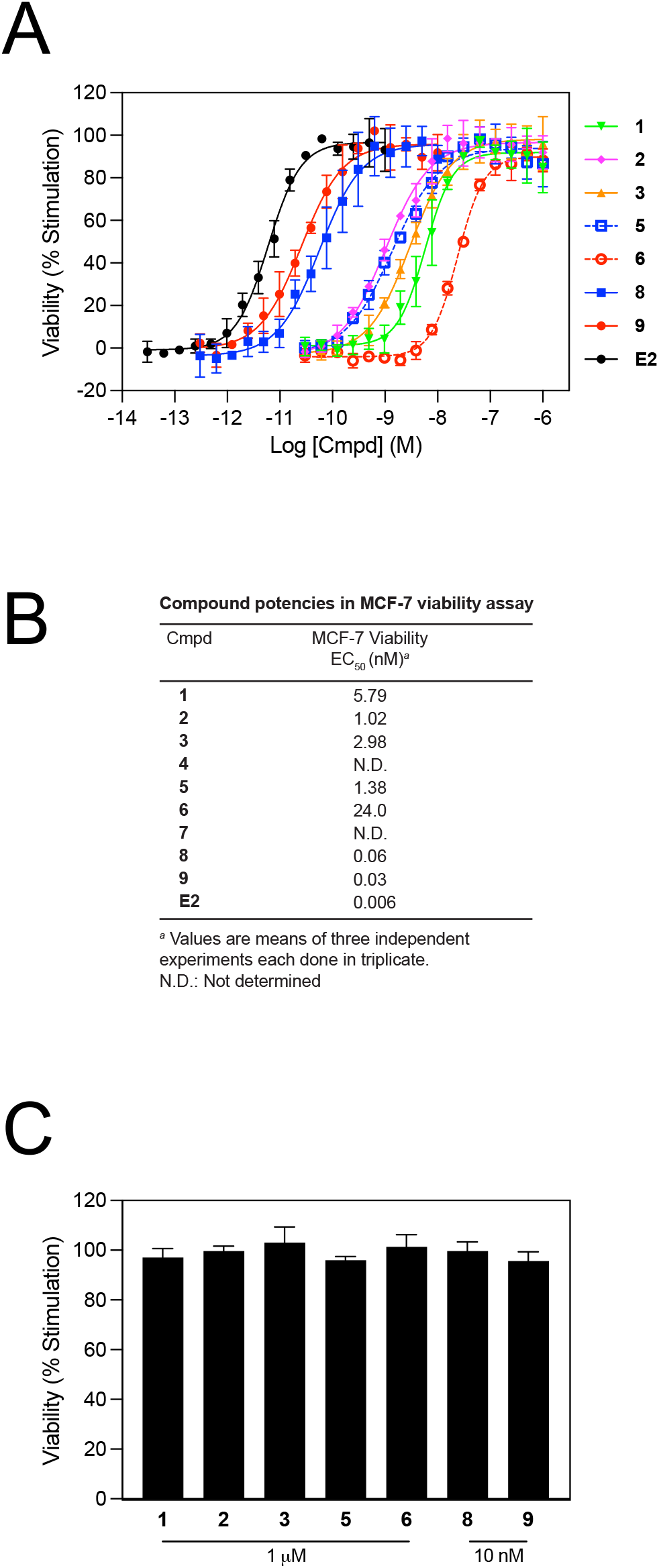
Effect of compounds **1-9** on cell viability of MCF-7 and T47D cells. (A) MCF-7 cells were treated with the indicated concentrations of compounds for 5 days and viability was assessed by CellTiter-Glo. (B) EC_50_ values of the dose-response curves shown in (A). (C) T47D cells were treated for 5 days with either 10 nM or 1μM of the indicated compounds and viability was assessed by CellTiter-Glo. Viability is shown in (A) and (C) as a percentage compared to positive (1 nM E2 in 0.1 % DMSO (v/v)) and negative (0.1 % DMSO (v/v)) controls. Values are the means ± standard deviations of 3 independent experiments each performed in triplicate. Dose-response for **E2** is provided as a reference.

Compound **1**, a known ERα-selective agonist [21], dose-dependently increased MCF-7 viability with an EC_50_ of 5.79 nM (**Fig. 3a and b**). Compounds **2**-**9** retained agonist activity, increasing cell viability with EC_50_ values ranging from 0.03-24.0 nM. These potencies largely correlated with their respective values in the ERα-LBD binding assay. The presence of the single *ortho*-fluoride in **2** resulted in a similar ∼5-fold increase in potency in both binding and viability assays relative to **1**. This functional benefit corresponds with an independent study wherein addition of this *ortho*-fluoride increased the functional potency of **1** in an ER-mediated transcription assay [19]. Interestingly, the carboxylic acid derivative **5**, with an ERα binding affinity comparable to **1**, exhibited a ∼4-fold improvement in potency over **1** in the viability assay. This discrepancy could be due to the fact that binding measured *in vitro* involves only the interaction between the ligand and the receptor, whereas cell-based assay activity additionally involves interactions between the ligand/receptor complex and other cellular components, such as coregulators and DNA. Thus, the relationship between *in vitro* binding affinities and cellular functional potencies may not be perfectly correlated due to other cellular components [24]. Although we generally observed a strong correlation between improved binding affinity and viability potency, **6** was an exception, showing ∼4-fold poorer potency in the viability assay compared to **1**. Compounds **8** and **9** were the most potent derivatives in the assay, exhibiting ∼100- and ∼200-fold improvements, respectively, in potency relative to **1**. The potency of **9** was remarkable, with only a 5-fold attenuation of EC_50_ value relative to the natural ligand of ERα (**E2**).

To assess whether the observed activities in our viability assay employing MCF-7 cells were cell line-specific, we similarly tested our compound series employing T47D cells. Like MCF-7 cells, T47D cells are also E2-sensitive, with E2 and other ERα agonists increasing their viability and proliferation. Based on the MCF-7 viability dose-response curves (**Fig. 3a**), using single-dose, saturating concentrations of compounds, we observed efficacies comparable to the E2 positive control across all tested compounds in T47D cells (**Fig. 3c**). Together these data show that our compound series is active in multiple E2-dependent cell lines.

### 2.4. Lead compound assessment in ERE-mediated transcription

In addition to overall viability and proliferation, we sought to determine the functional transcriptional activity of selected compounds in the series. Classical estrogen receptors bind to target sequences on DNA termed estrogen response elements (EREs) and subsequently modulate gene expression [25]. This mechanism is exploited in the MCF-7-E10 cell line, which stably expresses GFP under control of an ERE and induces GFP expression upon ER activation [21, 26]. This ER-mediated GFP expression can be quantified by flow cytometry. To this end, we used this cell line to assess the transcriptional potencies of several compounds in our series. Based on our ERα binding and viability data, **8** and **9** were identified as the lead compounds in our series, exhibiting nanomolar to subnanomolar potencies in both assays. We thus decided to assess the transcriptional potencies of these compounds employing MCF-7-E10 cells.

Compound **8** induced strong, dose-dependent GFP expression with an EC_50_ of 1.11 nM (**Figure 4**). This potency corresponds well with a previous report wherein **8** exhibited transcriptional potencies of 0.6 nM and 7.2 nM via ERα and ERβ, respectively, utilizing a different transcription assay [19]. This previously observed ∼10-fold preferential subtype potency cannot be tested in our assay due to endogenous expression of both ERα and ERβ in MCF-7-E10 cells. However, based on our binding data, which shows no preferential binding of **8** to either ER subtype, one might predict no transcriptional preference by **8**. This however remains speculative as binding affinity and functional potency do not always correlate, as exemplified by the binding/viability data of **5** and **6** in our study. Moreover, as noted above, binding measured using purified receptors only involves interactions between the ligand and the receptors while cell-based transcription involves additional interactions with other cellular components in complex with the receptor.

**Fig 4.**
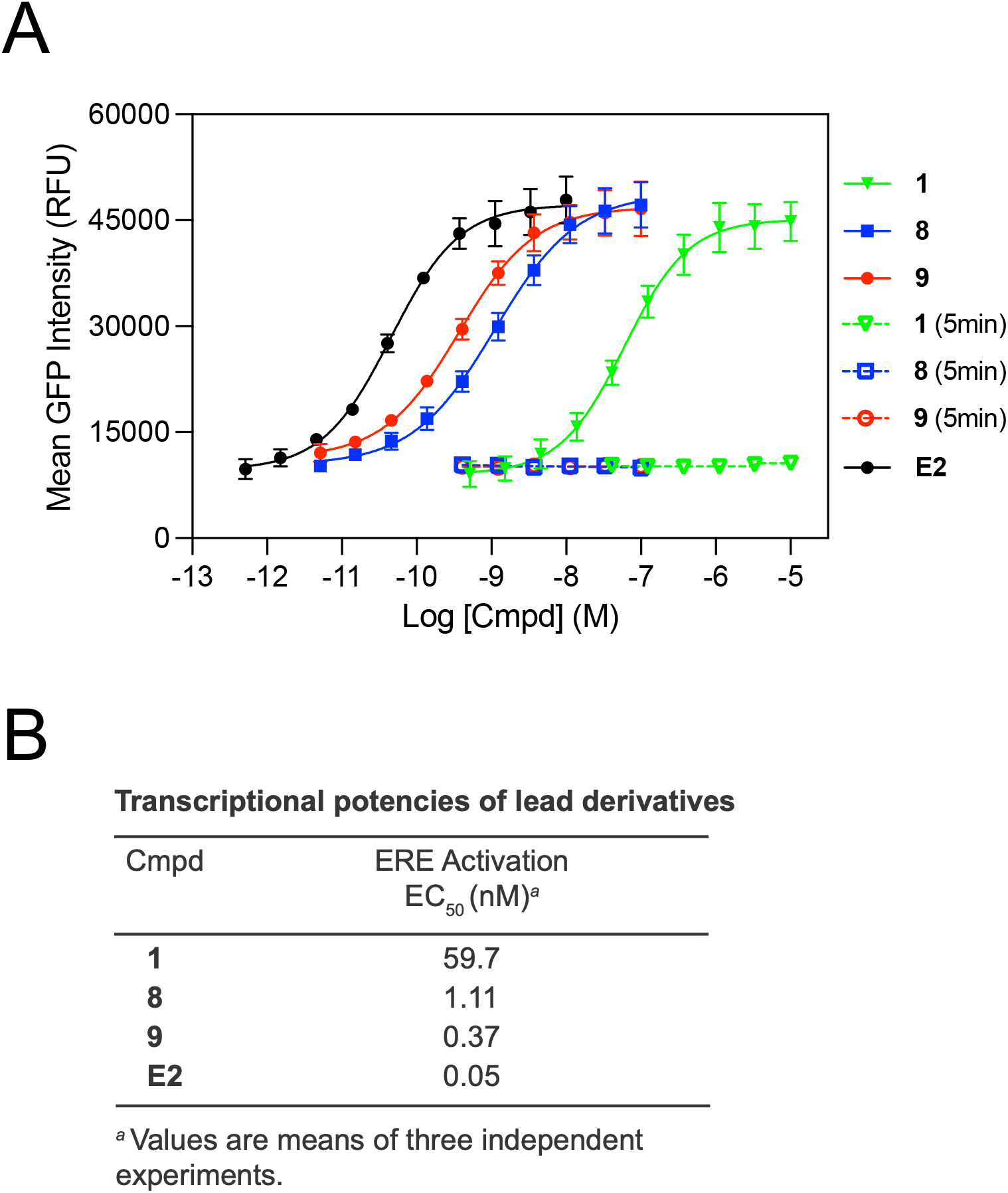
Transcriptional potencies of **1**, **8** and **9**. MCF-7-E10 cells were treated with the indicated concentrations of compounds for 24 h (*solid lines*) and GFP expression was measured by flow cytometry. Compound optical interference was assessed by treating MCF-7-E10 cells with the indicated concentrations of compounds for 5 min (*dashed lines*). (B) EC_50_ values of the dose-response curves shown in (A). Values are the means ± standard deviations of 3 independent experiments. RFU: relative fluorescence unit. Dose-response for **E2** is provided as a reference.

Derivative **9** outperformed **8** by 3-fold, achieving subnanomolar potency, with an EC_50_ of 0.37 nM. Such high potency is comparable to that of the ER-subtype selective compounds PPT (ERα EC_50_= ∼0.1 nM) and DPN (ERβ EC_50_= 0.85 nM) via their respective preferred receptor subtypes [5, 6]. Together, **8** and **9** represent a ∼50- and 160-fold improvement, respectively, over the EC_50_ of parent compound **1** (EC_50_= 59.7 nM, **Figure 4**) in this transcriptional assay, with **9** being only ∼7-fold less potent than **E2** (EC_50_= 0.05 nM). It is noteworthy that **9**, like all compounds in our series, is chiral and was studied as a racemate. Therefore, it is likely that an individual enantiomer of **9**, might exhibit a potency rivaling that of **E2**. Isolated enantiomers of **1** and **2** exhibit 1.8-3.7-fold improved potencies in a transcriptional assay compared to their racemates [19], thus supporting the feasibility of isolating enantiomers of **9** with further improved potency. This is an area we plan to examine in future work.

Lastly, to ensure that the tested compounds did not optically interfere in the fluorescence-based transcription assay, we treated the cells for 5 minutes with compound before measuring fluorescence. This short incubation allowed the compounds to associate with cells and bind ERs but not induce GFP transcription. We found no measurable increase in signal over our negative control (**Fig. 4a** *dashed lines*) indicating that the tested compounds were not optically interfering in the assay.

Taken together, we identified lead compound **9** as a highly potent ERα ligand harboring a binding affinity for the receptor rivaling that of **E2**. While **9** acts as an ERα agonist, its high affinity for the receptor could be exploited for potential therapeutic use in ERα-driven diseases. One such scenario could be its potential use as an ERα PROTAC for the treatment of ERα-positive breast cancers where ERα PROTACs, such as vepdegestrant, are currently being explored [27, 28]. In fact, a current PROTAC derived from the ERα agonist E2 exhibits anti-proliferative efficacy in cell models of ERα-positive breast cancers [29], implying that **9** could potentially be developed into an ERα PROTAC in the future.

## 3. Conclusion

We have reported the synthesis and evaluation of a series of oxabicyclo[3.3.1]nonene derivatives based on the scaffold of the ER agonist parent compound **1**. Through molecular modification of various regions of **1**, we have now identified two derivatives, **8** (previously discovered in [19]) and **9**, that exhibit low nanomolar to subnanomolar ERα binding affinities and potencies in ER-mediated functional assays, with **9** displaying potencies rivaling that of the natural ligand of ER. Furthermore, we discovered a key modification through reduction of the alkene group of the oxabicyclo[3.3.1]non-7-en-2-yl compound **8** to the saturated 3-oxabicyclo[3.3.1]nonan-2-yl derivative **9** that exhibited improved binding to ERα possibly attributed to improved van der Waals interactions with the hydrophobic binding pocket of the receptor. Finally, **8** and **9** exhibit low subtype selectivity of ERα over ERβ. The successful optimization described here, resulting in **9**, provides a solid foundation for future studies to address enantiomeric resolution and structural studies of **9** with the goal of potentially developing new ERα-targeted therapeutics.

## 4. Experimental section

### 4.1. Chemical synthesis

All compounds were synthesized in an efficient fume hood. Commercially available solvents and reagents were purchased and used without further purification. Compound **1** was prepared from diethyl 4-methylcyclohex-3-ene-1,1-dicarboxylate **10** and (4-methylcyclohex-3-ene-1,1-diyl)dimethanol **11** following the procedures previously described [21]. Column chromatography of compounds was performed using a Teledyne ISCO CombiFlash Rf+ Lumen chromatography system, with RediSep Silver disposable silica gel columns (sizes 12, 24, and 40 grams) and Supelco silica gel (pore size 60 Å, 40-63 μm particle size) for the stationary phase. ^1^H, ^13^C, and ^19^F NMR spectra were acquired using a Bruker Avance 500 spectrometer operating at 500 MHz, 125 MHz, and 470 MHz, respectively. The aforementioned spectra were collected at ambient temperatures (20 ± 2 °C). ^1^H, ^13^C, and ^19^F NMR spectra taken in CDCl_3_, acetone-d_6_, C_6_F_6_ or CD_3_OD were referred to tetramethylsilane (TMS). Mass spectra were obtained with assistance from the Mass Spectrometry Lab at the School of Chemical Sciences, University of Illinois Urbana-Champaign. Melting points of the compounds were measured using an Electrothermal-MEL-TEMP apparatus. A summary of multi-step syntheses and yields of analogs **2**-**9** are shown in **Table S1**.

#### General procedure for the synthesis of compound 1

##### 4-(5-(hydroxymethyl)-8-methyl-3-oxabicyclo[3.3.1]non-7-en-2-yl)phenol (1)

Detailed synthetic procedures and spectra related to **1** can be found in our previously published work [21].

##### General procedure for the Prins reaction for the synthesis of compounds 2 and 3

###### 2-fluoro-4-(5-(hydroxymethyl)-8-methyl-3-oxabicyclo[3.3.1]non-7-en-2-yl)phenol (2)

To a round bottom flask equipped with a magnetic stir bar under an N_2_ atmosphere was added compound **11** (118 mg, 0.75 mmol, 1.0 equiv.), 3-fluoro-4-hydroxybenzaldehyde (116.5 mg, 0.83 mmol, 1.1 equiv.), hafnium(IV) triflate (29.2 mg, 0.038 mmol, 0.05 equiv.) and acetonitrile (11.5 mL). The reaction flask was stirred at room temperature for 22 h. The reaction mixture was cooled to room temperature, diluted with ethyl acetate (25 mL) and transferred to a separatory funnel and then washed with saturated aqueous sodium bicarbonate solution (25 mL), then water (25 mL). The organic layer was dried over magnesium sulfate and concentrated by rotary evaporation. The crude mixture was purified by recrystallization from dichloromethane to afford the product **2** as a colorless solid. Yield: 67%. mp 146–148 °C. ^1^H NMR (CDCl_3_, 500 MHz) δ 7.07-7.04 (m, 1H), 6.96– 6.90 (m, 2H), 5.58-5.56 (m, 1H), 4.47 (s, 1H), 3.95 (dd, *J* = 10, 5 Hz, 1H), 3.61 (d, *J* = 10 Hz, 1H), 3.40 (s, 2H), 2.32-2.31 (m, 1H), 2.22-2.17 (m, 1H), 2.09-2.05 (m, 1H), 1.80 (dd, *J* = 15, 5 Hz, 1H), 1.66 (dt, *J* = 10, 5 Hz, 1H), 1.02 (q, *J* = 5 Hz, 3H). ^13^C NMR (CDCl_3_, 125 MHz) δ 150.9 (d, *J* = 236 Hz), 142.2 (d, *J* = 14 Hz), 135.1 (d, *J* = 6 Hz),, 132.95, 123.9, 121.6 (d, *J* = 4 Hz), 116.6 (d, *J* = 1 Hz), 112.8 (d, *J* = 19 Hz), 79.2, 77.7, 69.9, 42.6, 34.4, 33.8, 33.7, 24.0. ^19^F NMR (Acetone-*d*_6_, 470 MHz) δ −139.4 (**Figure S1**). FT-IR (neat) 3301, 2977, 1611, 1095, 1052 cm^−1^. Mass C_16_H_20_O_3_F^+^ [M+H]^+^ expected: 279.1396; found: 279.1398

###### 2,6-difluoro-4-(5-(hydroxymethyl)-8-methyl-3-oxabicyclo[3.3.1]non-7-en-2-yl)phenol (3)

The general procedure using 3,5-difluoro-4-hydroxybenzaldehyde (1.1 equiv.) was stirred at room temperature for 22 h to afford compound **3** as a colorless solid. Yield: 75%. mp 135–137°C. ^1^H NMR (CDCl_3_, 500 MHz) δ 6.87 (s, 1H), 6.86 (s, 1H), 5.58–5.56 (m, 1H), 4.43 (d, *J* = 5 Hz, 1H), 3.93 (dd, *J* = 10, 5 Hz, 1H), 3.60 (dd, *J* = 6, 1 Hz, 1H), 3.40 (s, 2H), 2.31–2.29 (m, 1H), 2.19–2.15 (m, 1H), 2.19-2.15 (m, 1H), 2.09-2.04 (m, 1H), 1.80 (dd, *J* = 14, 3 Hz, 1H), 1.80 (dd, *J* = 14, 3Hz 1H), 1.67 (dt, *J* = 10, 5 Hz, 1H), 1.04-1.03 (m, 3H). ^13^C NMR (CDCl_3_, 125 MHz) δ 152.5 (d, *J* = 6 Hz), 150.6 (d, *J* = 5 Hz), 134.0 (t, *J* = 7.5 Hz), 132.5, 131.2 (t, *J* = 16 Hz), 124.2, 108.8 (d, *J* = 6 Hz), 108.6 (d, *J* = 6 Hz), , 78.7, 77.7, 69.8, 42.5, 34.3, 33.8, 33.5, 23.9. ^19^F NMR (Acetone-*d*_6_, 470 MHz) δ −135.4 (**Figure S2**). FT-IR (neat) 3305, 2972, 1615, 1088, 1049 cm^−1^. Mass C_16_H_17_O_3_F_2_^−^ [M-H]^−^ expected: 295.1146; found: 295.1150 *4-(5-(hydroxymethyl)-8-methyl-3-oxabicyclo[3.3.1]non-7-en-2-yl)phenyl)boronic acid (**4**).* Sodium bicarbonate (20.75 mg, 0.247 mmol) was added to a solution of compound **13** (17 mg, 0.041 mmol) in methanol (10 mL) in a round bottom flask equipped with a magnetic stir bar. The reaction mixture was stirred at room temperature for 20 h then filtered and concentrated to give a crude residue that was recrystallized from ethyl acetate and hexane to afford the product **4** (10.6 mg) as a colorless amorphous solid. Yield: 90%. ^1^H NMR (Acetone-*d*_6_, 500 MHz,) δ 7.80 (d, *J* = 10 Hz, 2H), 7.33-7.31 (m, 2H), 5.50-5.48 (m, 1H), 4.56 (d, *J* = 2 Hz, 1H), 3.89 (dd, *J* = 10, 5, Hz, 1H), 3.61-3.59 (m, 1H), 3.31 (s, 1H), 2.41 (q, *J* = 5 Hz, 1H), 2.07–2.06 (m, 2H), 1.88 (dd, *J* = 10, 5 Hz, 1H), 1.63 (dt, *J* = 10, 5 Hz, 2H), 0.88 (m, 3H). ^13^C NMR (Acetone-*d*_6_,125 MHz,) δ 145.7, 134.5, 133.9, 125.6, 125.0, 80.7, 78.7, 43.9, 35.6, 34.9, 34.7, 30.3, 30.2, 24.3 (**Figure S3**). Mass C_16_H_20_BO_4_^−^ [M-H]^−^ expected: 287.1455; found: 287.1515.

##### General procedure for the Prins reaction for the synthesis of compounds 5 and 6

###### 4-(3-fluoro-4-hydroxyphenyl)-2,2,6-trimethyl-3-oxabicyclo[3.3.1]non-6-ene-1-carboxylic acid (5)

The general procedure using alcohol **15c** was stirred at room temperature overnight to yield a crude product that was recrystallized from ethanol and water to afford the compound **5** as colorless solid. Yield: 52%. mp 186–189 °C ^1^H NMR (Acetone-*d*_6_, 500 MHz) δ 7.06 (d, J = 10 Hz, 1H), 6.98 (d, *J* = 5 Hz, 1H), 6.91 (t, *J* = 10 Hz, 1H), 5.46-5.45 (m, 1H), 4.87 (d, *J* = 5 Hz, 1H), 2.80–2.75 (m, 2H), 2.64-2.61 (m, 1H), 2.38 (q, *J* = 5 Hz, 1H), 2.17-2.12 (m, 1H), 2.09-2.07 (m, 1H), 1.80 (dd, *J* = 15, 5 Hz, 1H), 1.49 (s, 3H), 1.36 (s, 3H), 0.94 (q, *J* = 5 Hz, 3H). ^13^C NMR (Acetone-*d*_6_, 125 MHz) δ 177.1, 152.1 (d, *J* = 237.5 Hz), 144.2 (d, *J* = 5 Hz), 133.2, 123.8, 122.4 (d, *J* = 4 Hz), 117.9 (d, *J* = 2.5 Hz), 114.1 (d, *J* = 20 Hz), 77.4, 73.0, 45.5, 43.3, 32.2, 31.6, 26.7, 23.9, 22.5. ^19^F NMR (Acetone-*d*_6_, 470 MHz) δ −139.4 (**Figure S4**). FT-IR (neat) 3450, 3310, 2970, 1708, 1602, 1098, 1053 cm^−1^. Mass C_18_H_20_O_4_F^−^ [M-H]^−^ expected: 319.1346; found: 319.1354.

###### 4-(3-fluoro-4-hydroxyphenyl)-2,2,6-trimethyl-3-oxabicyclo[3.3.1]non-6-ene-1-carbonitrile (6)

A mixture of **18**, 3-Fluoro-4-hydroxybenzaldehyde, and hafnium(IV) triflate in dry acetonitrile was heated to 50°C for 6 hours under an N_2_ atmosphere to yield a crude product that was purified by column chromatography (5–20% EA/Hex eluent) to afford the product **6** as a colorless solid. Yield: 70%. mp 177–180°C. ^1^H NMR (CDCl_3_, 500 MHz) δ 7.02 (d, J = 10 Hz, 1H), 6.94–6.90 (m, 2H), 5.44-5.43 (m, 1H), 4.82 (m, 1H), 2.83-2.78 (m, 1H), 2.62-2.59 (m, 1H), 2.50-2.45 (m, 1H), 2.29-2.27 (m, 1H), 2.07 (dd, *J* = 10, 5 Hz, 1H), 1.61 (s, 3H), 1.48 (s, 3H), 0.95 (s, 3H). ^13^C NMR (CDCl_3_, 125 MHz) δ 150.9 (d, *J* = 235 Hz), 142.4 (d, *J* = 15 Hz), 134.2 (d, J = 6 Hz), 124.1, 121.7 (d, J = 4 Hz), 121.3, 116.7 (d, J = 1 Hz), 113.0, 112.9, 76.4, 72.4 (d, J = 1 Hz), 40.3, 36.1, 31.9, 31.8, 26.5, 23.8, 21.9. ^19^F NMR (C_6_F_6_, 470 MHz) δ −144.2 (**Figure S5**). FT-IR (neat) 3300, 2977, 2350, 1612, 1095, 1059 cm^−1^. Mass C_18_H_19_O_2_NF^−^ [M-H]^−^ expected: 300.1400; found: 300.1406.

##### 4-(5-(aminomethyl)-4,4,8-trimethyl-3-oxabicyclo[3.3.1]non-7-en-2-yl)-2-fluorophenol (7)

To a cooled (0 °C) suspension of lithium aluminum hydride (22.68 mg, 0.6 mmol) in diethyl ether (2 mL) in a round bottom flask equipped with a magnetic stir bar and under an N_2_ atmosphere was added slowly over 5 min a solution of compound **6** (60 mg, 0.2 mmol,) in dry diethyl ether (8 mL). After the addition, the reaction was stirred at 37°C for 48 h. The reaction flask was cooled in an ice bath and quenched by sequential addition of water (23 μL), 10% aqueous NaOH solution (23 μL), and a second addition of water (69 μL) while stirring. The reaction mixture was filtered through celite to remove the aluminum salts and rinsed with dichloromethane (30 mL). The filtrate was concentrated by rotary evacuation and the crude product was purified by chromatography through a plug of silica gel using 10% methanol in DCM to afford compound **7** as a colorless solid. Yield: 55%. mp 84-86°C. ^1^H NMR (CD_3_OD, 500 MHz) δ 6.98–6.95 (m, 1H), 6.90-6.88)m, 1H), 6.82-6.79 (m, 1H), 5.50-5.48 (m, 1H), 4.77-4.76 (m, 1H), 2.60 (d, *J* = 15 Hz, 1H), 2.51 (d, *J* = 15 Hz, 1H), 2.40-2.36 (m, 1H), 2.29 (q, *J* = 5 Hz, 1H), 2.03-2.00 (m, 2H), 1.65 (dd, *J* = 15, 5 Hz, 1H), 1.36 (s, 3H), 1.27 (s, 3H), 0.93-0.92 (m, 3H). ^13^C NMR (CD_3_OD, 125 MHz) δ 152.9 (d, *J* = 237.5 Hz), 144.9 (d, *J* = 12.5 Hz),136.0 (d, *J* = 5 Hz), 133.8, 125.0, 122.8 (d, *J* = 2.5 Hz), 118.1, (d, *J* = 2.5 Hz), 114.6 (d, *J* = 20 Hz), 79.4, 73.8, 48.3, 44.2, 38.5, 31.4, 30.5, 26.2, 24.2, 22.0. ^19^F NMR (CD_3_OD, 470 MHz) δ −140.3 (**Figure S6**). FT-IR (neat) 3450, 3306, 2974, 1605, 1096, 1058 cm^−1^. Mass C_18_H_25_NO_2_F^+^ [M+H]^+^ expected: 306.1869; found: 306.1861.

##### 2-fluoro-4-(5-(hydroxymethyl)-4,4,8-trimethyl-3-oxabicyclo[3.3.1]non-7-en-2-yl)phenol (8)

To a cooled (0°C) suspension of lithium aluminum hydride (0.25 g, 6.458 mmol, 1.5 equiv.) in diethyl ether in a round bottom flask equipped with a magnetic stir bar and under an N_2_ atmosphere was added a solution of compound **16a** (1.50 g, 4.305 mmol, 1.0 equiv.) in dry diethyl ether at a dropwise rate. The reaction mixture was stirred at 0°C for 10 min, then warmed to room temperature and stirred for 2 h, then warmed to 40°C and stirred for 1 h. The reaction mixture was then cooled to 0 °C and ethyl acetate (10 mL, 25 equiv.) was added at a dropwise rate. The reaction mixture was treated successively with water (0.25 mL), 10% aqueous sodium hydroxide solution (0.25 mL), and a second addition of water (0.75 mL) while stirring. Magnesium sulfate was added and the reaction flask was equipped with a condenser and heated to reflux for 10 min to break up any remaining solids. The reaction mixture was filtered through celite and concentrated by rotary evaporation. The crude residue was purified by column chromatography (0–40% EA/Hex eluent) to yield the pure product **8** (1.098 g), as a white amorphous solid. Yield: 83%. ^1^H NMR (Acetone-d_6_, 500 MHz): δ 8.43 (br s, 1H), 7.05 (d, *J* = 15.0 Hz, 1H), 6.96 (d, *J* = 5.0 Hz, 1H), 6.89 (t, *J* = 10.0 Hz, 1H), 5.46 (s, 1H), 4.77 (s, 1H), 3.42 (d, *J* = 10.0 Hz, 1H), 3.27 (d, *J* = 15.0 Hz, 1H), 2.44 (m, 1H), 2.29 (s, 1H), 2.10 (m, 1H), 1.99 (d, *J* = 15.0 Hz, 1H), 1.58 (d, *J* = 10 Hz, 1H), 1.33 (s, 3H), 1.32 (s, 3H), 0.91 (s, 3H). ^13^C NMR (Acetone-d_6_, 125 MHz): δ 152.1 (d, J = 236 Hz), 144.0 (d, *J* = 12.5 Hz), 136.5 (d, *J* = 6 Hz), 132.7, 125.1, 122.4, 117.8 (d, *J* = 2.5 Hz), 114.3, (d, *J* = 20 Hz), 78.5, 73.1, 69.2, 43.6, 38.8, 31.5, 31.1, 27.1, 24.1, 21.9. ^19^F NMR (Acetone-d_6_, 470 MHz): δ −139.4 (**Figure S7**). IR (neat), cm^−1^: 3337, 2932, 1518, 1437, 1206, 1151. HRMS (EI+) m/z: [M]^+^ C_18_H_23_O_3_F^+^ expected: 306.1631; found: 306.1638.

##### 2-fluoro-4-(5-(hydroxymethyl)-4,4,8-trimethyl-3-oxabicyclo[3.3.1]nonan-2-yl)phenol (9)

In a 250-mL glass hydrogenation vessel, compound **8** (0.45 g, 1.556 mmol, 1.0 equiv.) was dissolved in ethanol (52 mL, 0.03 M) at room temperature. To this was added 10% w/w palladium on carbon catalyst (0.662 g, 0.622 mmol, 40 mol%). The vessel was then attached to a shaker-type hydrogenator apparatus and shaken under 32 psi hydrogen gas at room temperature for 5 h. The reaction mixture was removed from the hydrogenator and filtered through celite and rinsed with ethyl acetate (approximately 20 mL total). The combined organic material was concentrated using a rotary evaporator. The crude residue was recrystallized from hot DCM to afford the product **9** (0.208 g) as a white solid. Yield: 52%. The overall yield of **9** over five steps with a final purification by recrystallization was 21 %. mp 154.0°C. ^1^H NMR (Acetone-d_6_, 500 MHz): δ 8.34 (s, 1H), 7.16 (d, *J* = 15.0 Hz, 1H), 7.04 (d, *J* = 5.0 Hz, 1H), 6.91 (t, *J* = 5.0 Hz, 1H), 4.90 (s, 1H), 3.64-3.60 (m, 1H), 3.40 (dd, *J* = 15.0, 5.0 Hz, 1H), 3.21 (dd, *J* = 10.0, 5.0 Hz, 1H), 2.17 (d, *J* = 15.0 Hz, 1H), 2.10–2.01 (m, 4H), 1.66 (br s, 1H), 1.43–1.31 (m, 3H), 1.40 (s, 3H), 1.29 (s, 3H), 0.34 (d, *J* = 10.0 Hz, 3H). ^13^C NMR (Acetone-d_6_, 125 MHz): δ 152.0 (d, J = 236 Hz), 143.6 (d, *J* = 13.75 Hz), 137.4 (d, *J* = 6.25 Hz), 122.2 (d, *J* = 3.75 Hz), 117.8 (d, *J* = 2.5 Hz), 113.8 (d, *J* = 18.75 Hz), 78.7, 74.4, 70.0, 41.3, 39.2, 38.4, 37.1, 32.4, 31.0, 26.5, 22.4, 21.7. ^19^F NMR (Acetone-*d*_6_, 470 MHz) δ −140.5 (**Figure S8**). IR (neat) cm^−1^: 3351, 2946, 1599, 1518, 1453, 1379. HRMS (EI+) m/z: [M]^+^ C_18_H_25_O_3_F^+^ expected: 308.1788; found: 308.1782.

##### 8-(4-(5-(hydroxymethyl)-8-methyl-3-oxabicyclo[3.3.1]non-7-en-2-yl)phenyl)-4-methyldihydro-4l^4^,8l^4^-[1,3,2]oxazaborolo[2,3-b][1,3,2]oxazaborole-2,6(3H,5H)-dione (13)

A mixture of alcohol **11** (25 mg, 0.16 mmol), and 4-formylphenylboronic acid MIDA ester **12** was heated to 40^°^C for 19 h to afford product **13** as a colorless solid. Yield: 61%. mp 228–230°C. Product **13** was used in the next step without further purification. ^1^H NMR (Acetone-*d*_6_, 500 MHz,) δ 7.45, *J* = 8.1 Hz, 2H), 7.33 (d, *J* = 7.75 Hz, 2H), 5.49–5.47 (m, 1H), 4.57 (d, *J* = 1.6 Hz, 1H), 4.33 (d, *J* = 17.0 Hz, 2H), 4.11 (dd, *J* = 17.0, 2.7 Hz, 2H), 3.89 (dd, *J* = 10.8, 2.6 Hz, 1H), 3.69 (t, *J* = 4.25 Hz, 1H), 3.30 (d, *J* = 10.8 Hz, 2H), 2.80 (s, 1H), 2.68 (s, 3H), 2.41–2.39 (m, 1H), 2.11–2.07 (m, 2H), 1.88 (dd, *J* = 11.8, 2.9 Hz, 1H), 1.63 (dt, *J* = 11.7, 3.05 Hz 1H), 0.87 (td, *J* = 2.2, 1.4 Hz, 3H). ^13^C NMR (Acetone-*d*_6_,125 MHz,) δ 169.30, 144.46, 133.81, 132.70, 125.87, 124.90, 80.56, 78.58, 69.74, 62.70, 48.22, 43.88, 35.49, 34.81, 34.60, 24.25.

##### General procedure for Diels-Alder reaction with acetoacetic esters for the synthesis of compounds 14a and 14b

###### Ethyl 1-acetyl-4-methylcyclohex-3-ene-1-carboxylate (14a)

To a thick-walled glass reaction flask equipped with a magnetic stir bar was added ethyl acetoacetate (1.50 g, 11.53 mmol, 1.0 equiv.), paraformaldehyde (0.520 g, 17.30 mmol, 1.5 equiv.), isoprene (3.95 g, 57.65 mmol, 5.0 equiv.), THF (46 mL, 0.25 M), and zinc(II) chloride (0.314 g, 2.306 mmol, 20 mol%). The system was purged with N_2_ and the flask sealed with a screw-top cap fitted with an O-ring and Teflon tape. The reaction flask was heated in an oil bath at 70°C while stirring for 20 h. The reaction mixture was cooled to room temperature and then transferred to a round bottom flask and concentrated by rotary evaporation to a viscous oil. The crude residue was purified by column chromatography (0–10% EA/Hex eluent) to yield the product **14a** (1.70 g) as a colorless oil. Yield: 70%. ^1^H NMR (CDCl_3_, 500 MHz): δ 5.38–5.34 (m, 1H), 4.22–4.13 (m, 2H), 2.53 (d, *J* = 17.25 Hz, 1H), 2.41 (d, *J* = 11.45 Hz, 1H), 2.17 (s, 3H), 2.15–1.91 (m, 4H), 1.62 (s, 3H), 1.24 (t, *J* = 5.0 Hz, 3H). ^13^C NMR (CDCl_3_, 125 MHz): δ 205.4, 172.3, 133.7, 118.1, 61.5, 59.1, 30.1, 27.30, 27.27, 25.96, 23.4, 14.2. IR (neat) cm^−1^: 2931, 1713, 1446, 1356, 1289, 1207, 1152. HRMS (ES+) m/z: [M + H]^+^ Calcd for C_12_H_19_O_3_^+^ 211.1334; found 211.1335. ***Note***:

The product isolated consists of an 87:13 mixture of **14a** and the regioisomer, with the ratio determined by comparison of integration values between the ^1^H NMR signals of **14a** at δ 1.62 and 2.53 with the corresponding signals of the minor regioisomer at δ 1.70 and 2.34–2.27.

###### Tert-butyl 1-acetyl-4-methylcyclohex-3-ene-1-carboxylate (14b)

From *tert*-butyl acetoacetate the product **14b** was obtained as a colorless oil. Yield: 85%. ^1^H NMR (CDCl_3_, 500 MHz): δ 5.38–5.34 (m, 1H), 2.48 (d, *J* = 17.3 Hz, 1H), 2.34 (d, *J* = 16.6 Hz, 1H), 2.29– 2.20 (m, 1H), 2.17 (s, 3H), 2.13–1.90 (m, 3H), 1.62 (s, 3H), 1.44 (s, 9H). ^13^C NMR (CDCl_3_, 125 MHz) δ 205.4, 172.3, 133.7, 118.1, 61.5, 59.1, 30.1, 27.30, 27.27, 25.96, 23.4, 14.2. IR (neat) cm^−1^: 2931, 1713, 1446, 1356, 1289, 1207, 1152. ***Note:*** The product isolated consists of an 80:20 mixture of **14b** and the regioisomer, with the ratio determined by comparison of integration values between the ^1^H NMR signals of **14b** at δ 1.62 and the corresponding signal of the minor regioisomer at δ 1.70.

##### General procedure for Grignard addition for the synthesis of compounds 15a and 15b

###### Ethyl 1-(2-hydroxypropan-2-yl)-4-methylcyclohex-3-ene-1-carboxylate (15a)

In a round bottom flask equipped with a magnetic stir bar under an N_2_ atmosphere, compound **14a** (1.55 g, 7.371 mmol, 1.0 equiv.) was dissolved in THF (37 mL) and cooled to 0 °C. At this temperature, methylmagnesium iodide (3.0 M in diethyl ether, 4.9 mL, 2.0 equiv.) was added dropwise, while stirring. The reaction mixture was stirred at 0°C for 5 h. The reaction was quenched with addition of saturated aqueous ammonium chloride solution (20 mL), then transferred into a separatory funnel. The aqueous layer was drained, and the organic layer was washed with water (20 mL). The organic layer was dried over magnesium sulfate, filtered and concentrated using a rotary evaporator. The crude residue was purified by column chromatography (0–20% EA/Hex eluent) to yield the pure product **15a** as a colorless oil (1.36 g). Yield: 82%. ^1^H NMR (CDCl_3_, 500 MHz): δ 5.38–5.36 (m, 1H), 4.21–4.16 (m, 2H), 2.92 (s, 1H), 2.55–2.50 (m, 1H), 2.22–2.17 (m, 1H), 2.12–2.02 (m, 2H), 1.96–1.92 (m, 1H), 1.63–1.59 (m, 1H), 1.61 (s, 3H), 1.26 (t, *J* = 5.0 Hz, 3H), 1.19 (s, 3H), 1.17 (s, 3H). ^13^C NMR (CDCl_3_, 125 MHz): δ 176.3, 133.4, 119.8, 73.7, 60.9, 53.3, 29.0, 28.5, 26.1, 25.8, 25.7, 23.3, 14.3. IR (neat) cm^−1^: 3502, 2974, 1714, 1445, 1374, 1289, 1204, 1150. HRMS (ES+) m/z: [M + Na]^+^ Calcd for C_13_H_22_O_3_Na^+^ 249.1467; found 249.1469. ***Note:*** The product isolated consists of an 85:15 mixture of **15a** and the regioisomer with the ratio determined by the comparison of integration values between the ^1^H NMR signals of **15a** at δ 5.38–5.36 with the corresponding signal of the minor regioisomer at δ 5.33–5.32.

###### Tert-butyl 1-(2-hydroxypropan-2-yl)-4-methylcyclohex-3-ene-1-carboxylate (15b)

The general procedure for compound **14b** with methylmagnesium iodide (2.0 equiv.) afforded the product **15b** as a colorless oil. Yield: 87%. ^1^H NMR (CDCl_3_, 300 MHz) δ 5.38 (m, 1H), 3.25 (s, 1H), 2.47–2.42 (m, 1H), 2.28–2.13 (m, 1H), 2.07–2.01 (m, 2H), 1.97– 1.92 (m, H), 1.64–1.55 (m, 1H), 1.62 (s, 3H), 1.44 (s, 9H), 1.19 (s, 3H), 1.18 (s, 3H). ^13^C NMR (CDCl_3_, 125 MHz) δ 175.70, 133.14, 120.03, 81.67, 73.65, 53.52, 29.21, 28.59, 28.08, 26.18, 25.95, 25.48, 23.24. FT-IR (neat) 3400, 1733, 1468, 1226 cm^−1^. ***Note:*** The product isolated consists of an 82:18 mixture of **15b** and the regioisomer with the ratio determined by the comparison of integration values between the ^1^H NMR signal of **15b** at δ 5.38 with the corresponding signal of the minor regioisomer at δ 5.32.

##### 1-(2-hydroxypropan-2-yl)-4-methylcyclohex-3-ene-1-carboxylic acid (15c)

To a solution of compound **15b** (50 mg, 0.196 mmol) in DCM (3 mL) in a round bottom flask equipped with a magnetic stir bar and cooled to 0°C was added dropwise trifluoroacetic acid (1.2 mL). The reaction mixture was stirred for 1 h then quenched with water (25 mL) and transferred to a separatory funnel. The mixture was extracted with ethyl acetate (3 x 50 mL). The combined organic layers were dried over magnesium sulfate, filtered and concentrated by rotary evaporation. The crude material was purified by column chromatography (10% EA/Hex eluent) to give **15c** (29.5 mg) as a colorless oil. Yield: 76%. ^1^H NMR (CDCl_3_, 500 MHz) δ 5.39 (m, 1H), 2.54–2.46 (m, 1H), 2.20–2.06 (m, 4H), 2.06–1.93 (m, 4H), 1.84–1.77 (m, 2H), 1.71–1.59 (m, 4H), 1.27 (s, 3H), 1.25 (s, 3H). FT-IR (neat) 3450, 3288, 1710, 1470, 1218 cm^−1^. ^13^C NMR (CD_3_OD, 125 MHz) δ 178.98, 133.96, 121.19, 73.97, 54.24, 29.71, 29.39, 26.37, 26.30, 26.06, 23.37.

##### Ethyl-4-(3-fluoro-4-hydroxyphenyl)-2,2,6-trimethyl-3-oxabicyclo[3.3.1]non-6-ene-1-carboxylate (16a)

A mixture of alcohol **15a**, 3-Fluoro-4-hydroxybenzaldehyde, and hafnium(IV) triflate in dichloromethane was stirred at room temperature for 24 h to give the crude product that was purified by recrystallization from ethyl acetate and hexane to afford the product **16a** as a yellow crystalline solid. Yield: 83%. mp 115.0°C. ^1^H NMR (CDCl_3_, 500 MHz): δ 7.07 (d, *J* = 15.0 Hz, 1H), 6.93 (dt, *J* = 15.0, 10.0 Hz, 2H), 5.46 (br s, 1H), 5.32–5.11 (br m, 1H) 4.76 (s, 1H), 4.19–4.08 (m, 2H), 2.79 (d, *J* = 15.0 Hz, 1H), 2.59 (d, *J* = 15.0 Hz, 1H), 2.30 (s, 1H), 2.11, (d, *J* = 20.0 Hz, 1H), 1.83 (d, *J* = 10.0 Hz, 1H), 1.48 (s, 3H), 1.32 (s, 3H), 1.27 (t, *J* = 5.0 Hz, 3H), 0.95 (s, 3H). ^13^C NMR (CDCl_3_, 125 MHz): δ 175.9, 151.1 (d, *J* = 236 Hz), 142.3 (d, *J* = 15 Hz), 135.6 (d, *J* = 5 Hz), 132.4, 123.2, 121.8 (d, J = 3.75 Hz), 116.7 (d, *J* = 2.5 Hz), 113.1 (d, *J* = 18.75 Hz), 77.1, 72.7, 60.7, 45.3, 42.5, 31.4, 30.7, 26.3, 23.8, 22.2, 14.2 ^19^F NMR (CDCl_3_, 470 MHz): δ −141.1. IR (neat) cm^−1^: 3397, 2978, 2906, 1718, 1518, 1437, 1151. HRMS (EI+) m/z: [M]^+^ C_20_H_25_O_4_F^+^ expected: 348.1737; found: 348.1725.

##### Ethyl 1-cyano-4-methylcyclohex-3-ene-1-carboxylate (17)

To a thick-walled glass reaction flask equipped with a magnetic stir bar was added ethyl 2-cyanoacrylate (0.40 mL, 3.39 mmol), isoprene (1.017 mL, 10.17 mmol), THF (17 mL), and zinc(II) chloride (46.4 mg, 0.339 mmol). The system was purged with N_2_ and the flask sealed with a screw-top cap fitted with an O-ring and Teflon tape. The reaction mixture was stirred at room temperature for 4 hours. The reaction mixture was filtered through celite with diethyl ether and the filtrate was concentrated using a rotary evaporator to give the product **17** as a colorless oil. Yield: 90%. ^1^H NMR (CDCl_3_, 300 MHz) δ 5.37 (m, 1H), 4.28 (q, *J* = 8 Hz, 2H), 2.56 (bs, 2H), 2.34–1.96 (m, 4H), 1.72 (s, 3H), 1.34 (t, *J* = 6 Hz, 3H). ^13^C NMR (CDCl_3_, 125 MHz) δ 169.10, 134.09, 119.32, 115.94, 62.75, 42.24, 32.52, 29.31, 26.88, 23.20, 13.96. FT-IR (neat) cm^−1^: 2210, 1745, 1476. See also [30].

##### General procedure for Grignard addition for the synthesis of compound 18

###### 1-(2-hydroxypropan-2-yl)-4-methylcyclohex-3-ene-1-carbonitrile (18)

The general procedure with compound **17** using methylmagnesium iodide (5.0 equiv.) afforded the product **18** as a colorless oil. Yield: 83%. ^1^H NMR (CDCl_3_, 300 MHz) δ 5.37 (m, 1H), 2.40– 2.20 (m, 3H), 2.15–2.03 (m, 2H), 1.72 (dq, *J* = 3.1, 1.6 Hz, 3H), 1.69–1.57 (m, 2H), 1.40 (d, *J* = 3.0 Hz, 3H), 1.35 (s, 3H). ^13^C NMR (CDCl_3_, 125 MHz) δ 134.36, 122.84, 117.33, 73.07, 46.35, 30.33, 28.26, 26.02, 25.88, 25.04, 23.13. FT-IR (neat) 2355, 2205, 1477, 1223 cm^−1^.

### 4.2. Cell culture

MCF-7 cells (ATCC, #HTB-22) were cultured in Dulbelcco’s Modified Eagle Medium (DMEM, Corning, #10-013-CV) without sodium pyruvate supplemented with 10 % fetal bovine serum (FBS, Corning, #35-016-CV), 2 mM L-Glutamine, 100 units/mL penicillin and 100 μg/mL streptomycin (Corning, #30-009-CI). MCF-7-E10 cells [26] were cultured as described above with the addition of 1 mg/mL G418 (Gibco, #11811-023). T47D cells (ATCC, #HTB-133) were cultured in RPMI 1640 (Corning, #10-040-CV) supplemented with 10 % FBS, 2 mM L-Glutamine, 100 units/mL penicillin and 100 μg/mL streptomycin. All cell lines were maintained in a humidified incubator at 37°C in 5 % CO_2_.

### 4.3. TR-FRET ERα/β-LBD ligand-binding assay

Binding assays for ERα/β-LBD were performed by the SelectScreen™ Biochemical Nuclear Receptor Profiling Service group (ThermoFisher Scientific) using the LanthaScreen™ TR-FRET Competitive Binding Assay for ERα and ERβ (ThermoFisher Scientific, #A15887 and A15895). The fluorescent E2 analog structure is not disclosed by the manufacturer (proprietary). Dose-response curves were assessed using 10-point, 3-fold dilutions with 1 μM or 10 μM as the upper limit. For highly potent compounds, an upper limit of 100 nM was used to achieve more accurate IC_50_ values. Fluorescent E2 displacement values were normalized to negative (1 % DMSO (v/v)) and positive (1 μM E2 in 1 % DMSO (v/v)) controls.

### 4.4. Cell viability assay

MCF-7 cells were plated (2,500 cells/well) in the center 60 wells of white, clear-bottom 96-well plates (Greiner Bio-One, #655098) in 100 μL phenol red-free DMEM/F-12 (Corning, #16-405-CV) supplemented with 10 % charcoal-stripped FBS (Sigma, #F6765), 2 mM L-Glutamine, 100 units/mL penicillin and 100 μg/mL streptomycin. The outer 24 wells were filled with medium without cells. Cells were maintained in a humidified incubator at 37°C in 5 % CO_2_. Twenty-four hours post-seeding, the medium was replaced with 100 μL medium containing the appropriate concentration of compound in the presence of 0.1 % DMSO (v/v) (Invitrogen, #D12345) final for 120 h at 37°C in 5 % CO_2_. The final volume per well was 100 μL with triplicate sample wells employed. The stimulation medium was refreshed after the first 72 h of stimulation. Compound dose responses were assessed using 16-point doubling dilutions with 1 nM, 1 μM or 10 μM as the upper limit. Plates were removed from the incubator for the last 30 min of stimulation and allowed to come to room temperature. Viability was subsequently assessed using the CellTiter-Glo luminescence assay (Promega, #G7571) (100 μL/well) according to the manufacturer’s instructions. Luminescence was measured on a Victor Nivo Multimode Plate Reader (PerkinElmer) using a 1 s integration time. Viability was expressed as a percentage compared to positive (1 nM E2 in 0.1 % DMSO (v/v)) and negative (0.1 % DMSO (v/v)) controls located on the same plate and defined as % stimulation = ((read value - negative control average)/(positive control average - negative control average)) × 100. Dose-response curves were fitted using a nonlinear regression variable slope (four parameters) in Prism 10 (GraphPad Software Inc.). Due to a stocking issue, one biological replicate of each MCF-7 plate was assayed using charcoal-stripped FBS purchased from Corning (#35-072-CV). For T47D cells (2,500 cells/well), viability was assessed as described above, but cells were treated for 120 h with single dose compounds (10 nM or 1 μM) in triplicate in the presence of 0.1 % DMSO (v/v) diluted in phenol red-free RPMI 1640 (Corning, #17-105-CV) supplemented with 10 % charcoal-stripped FBS (Corning, #35-072-CV), 2 mM L-Glutamine, 100 units/mL penicillin and 100 μg/mL streptomycin. Data was analyzed and as described above.

### 4.5. ERE-GFP reporter transcription assay

At ∼ 70 % culture confluency, MCF-7-E10 cells were harvested and plated in 6-well plates (Corning, #3516) (150,000 cells/well) in 2 mL phenol red-free DMEM/F-12 supplemented with 5 % charcoal-stripped FBS (Gibco, #12676-029), 2 mM L-Glutamine, 100 units/mL penicillin and 100 μg/mL streptomycin and allowed to acclimate for 5 days at 37°C in 5 % CO_2_. The medium was refreshed once on days 3 and 4 post-seeding. After 5 days, cells were treated for 24 h with 10-point, 3-fold dilutions of compounds in medium in the presence of 0.1 % DMSO (v/v) with 10 nM, 100 nM or 1 μM as the upper limit. After 24 h, the cells were rinsed twice with PBS (Corning, #21-040-CV) and detached for 5 min at 37°C with 0.5 mL/well phenol red-free trypsin (Gibco, #15400-054). Trypsin was deactivated by addition of 0.5 mL/well ice-cold medium (as described above) and cells were centrifuged for 5 min at 420 *x g* at 4°C. The pellet was reconstituted in 800 μL ice-cold PBS, filtered into Falcon tubes (Corning, #352235) and subsequently analyzed for GFP fluorescence on an Accuri C6 Plus flow cytometer (BD Biosciences) with excitation at 488 nm and emission at 525 nm. Ten thousand events (singlet-gated) were collected per sample. Positive (10 nM E2 in 0.1 % DMSO (v/v)) and negative (0.1 % DMSO (v/v)) controls were freshly prepared in duplicate each day alongside each dose-response samples. Mean fluorescence measurements were analyzed on the Accuri C6 Plus software (BD Biosciences) and are reported as relative fluorescence units (RFUs). Dose-response curves were fitted using a nonlinear regression variable slope (four parameters) in Prism 10.

To assess potential optical interference of the compounds, cells were treated for 5 min with 6-point, 3-fold dilutions of compound in the presence of 0.1 % DMSO (v/v) with 100 nM or 10 μM as the upper limit. The samples were further processed as described above.

## Acknowledgements

This research supported by NIH R01 grants CA163890 and CA194496 (E.R.P.), the Cowboys for Cancer Research Foundation and the Center of Biomedical Research Excellence in Autophagy, Inflammation and Metabolism (P20 GM121176). This research was partially supported by UNM Comprehensive Cancer Center Support Grant NCI P30CA118100 and the Flow Cytometry shared resource. We would also like to acknowledge the dissertation work of Dr. Travis A. Saxon that provided the basis for the syntheses described in the current study which is available through the online repository ProQuest [31].

## Abbreviations

C_6_F_6_: hexafluorobenzene
CDCl_3_: deuterochloroform
CD_3_OD: methanol-d_4_
DCM: dichloromethane
DPN: diarylpropionitrile
EA: ethyl acetate
Hex: hexanes
MeOH: methanol
MIDA: N-methyliminodiacetic acid
PPT: propylpyrazole triol
TFA: trifluoroacetic acid
THF: tetrahydrofuran

## Scheme Legends

**Scheme 1.**
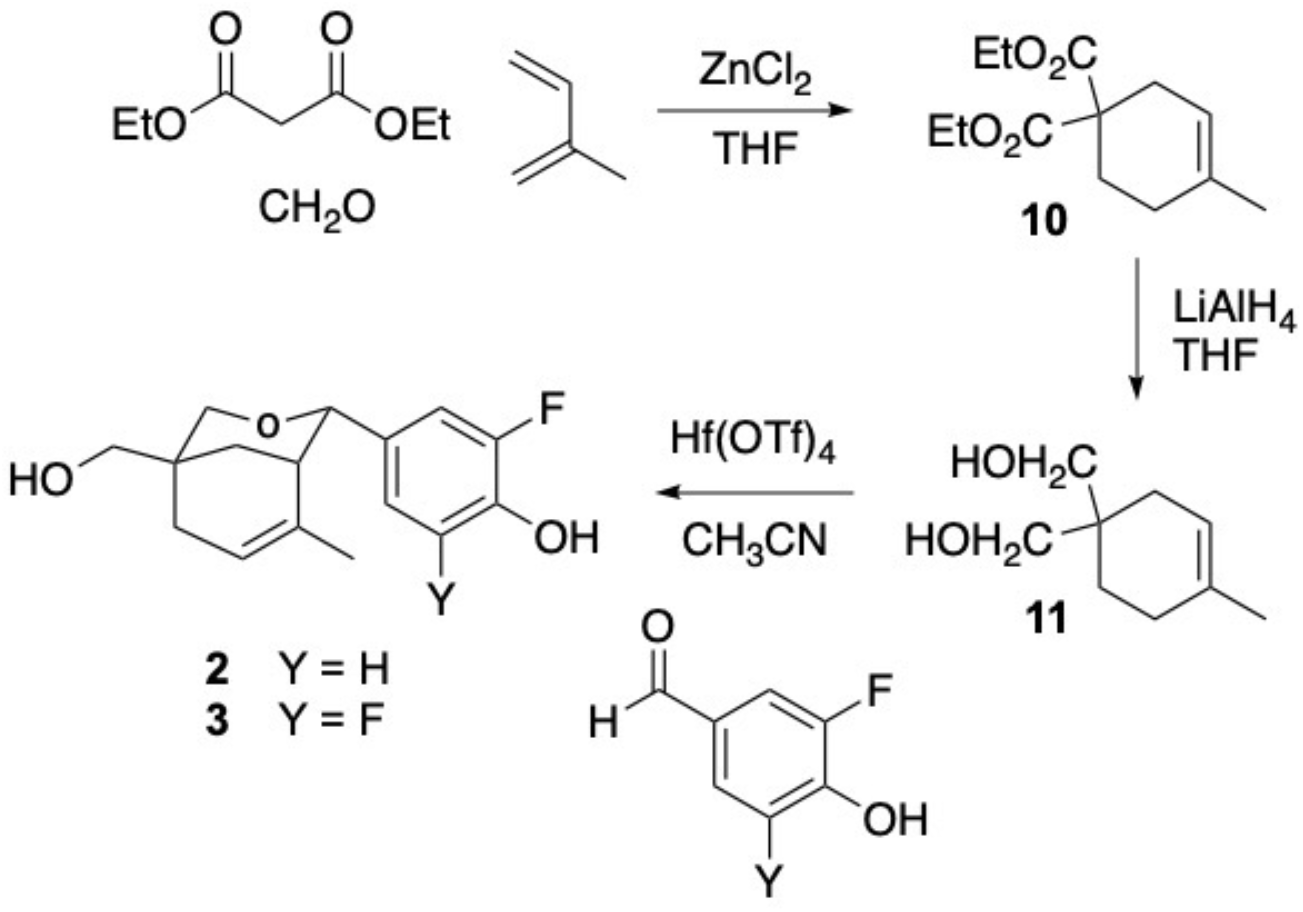
General procedure for the synthesis of compounds **2** and **3**.

**Scheme 2.**
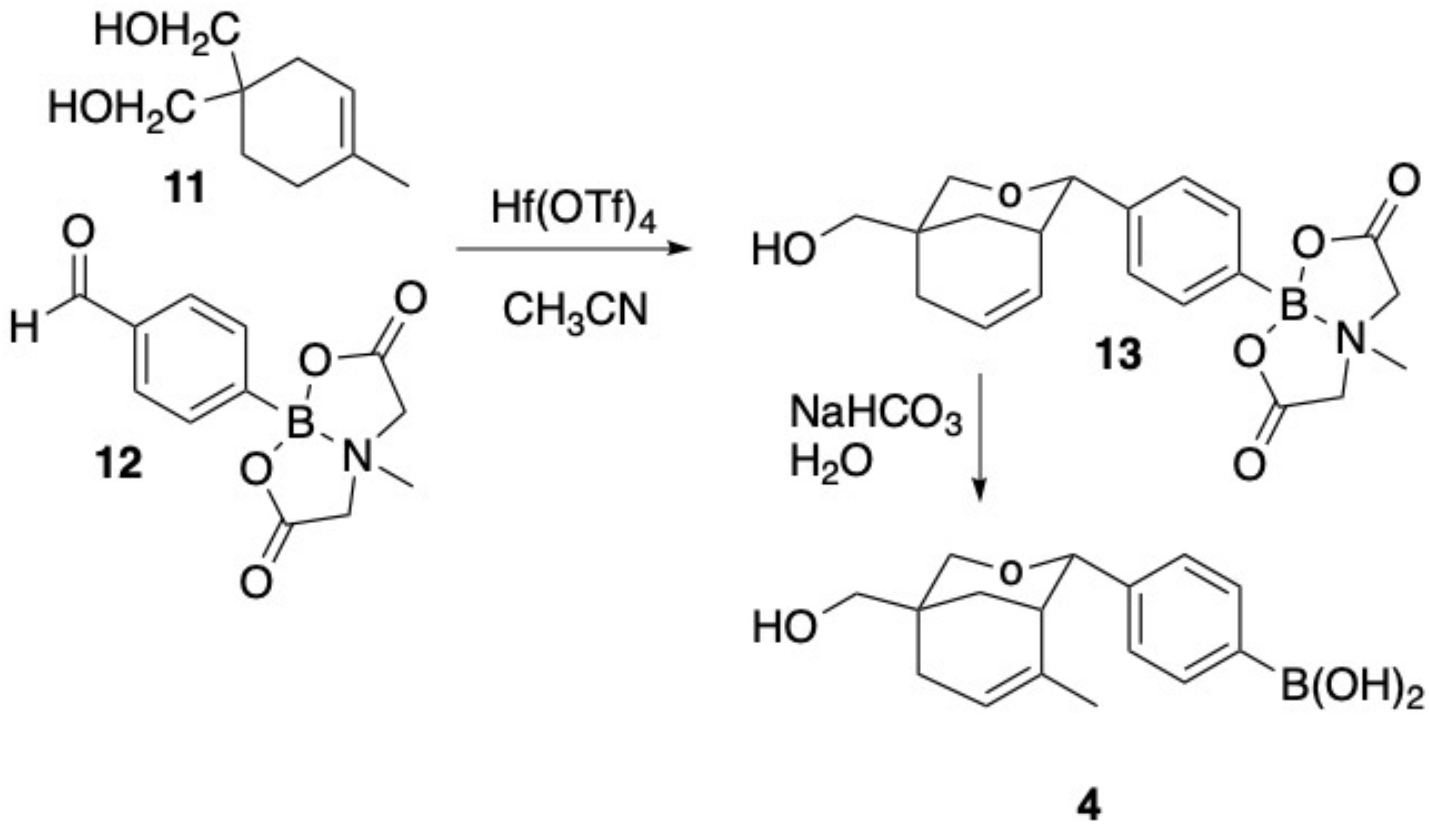
General procedure for the synthesis of compound **4**.

**Scheme 3.**
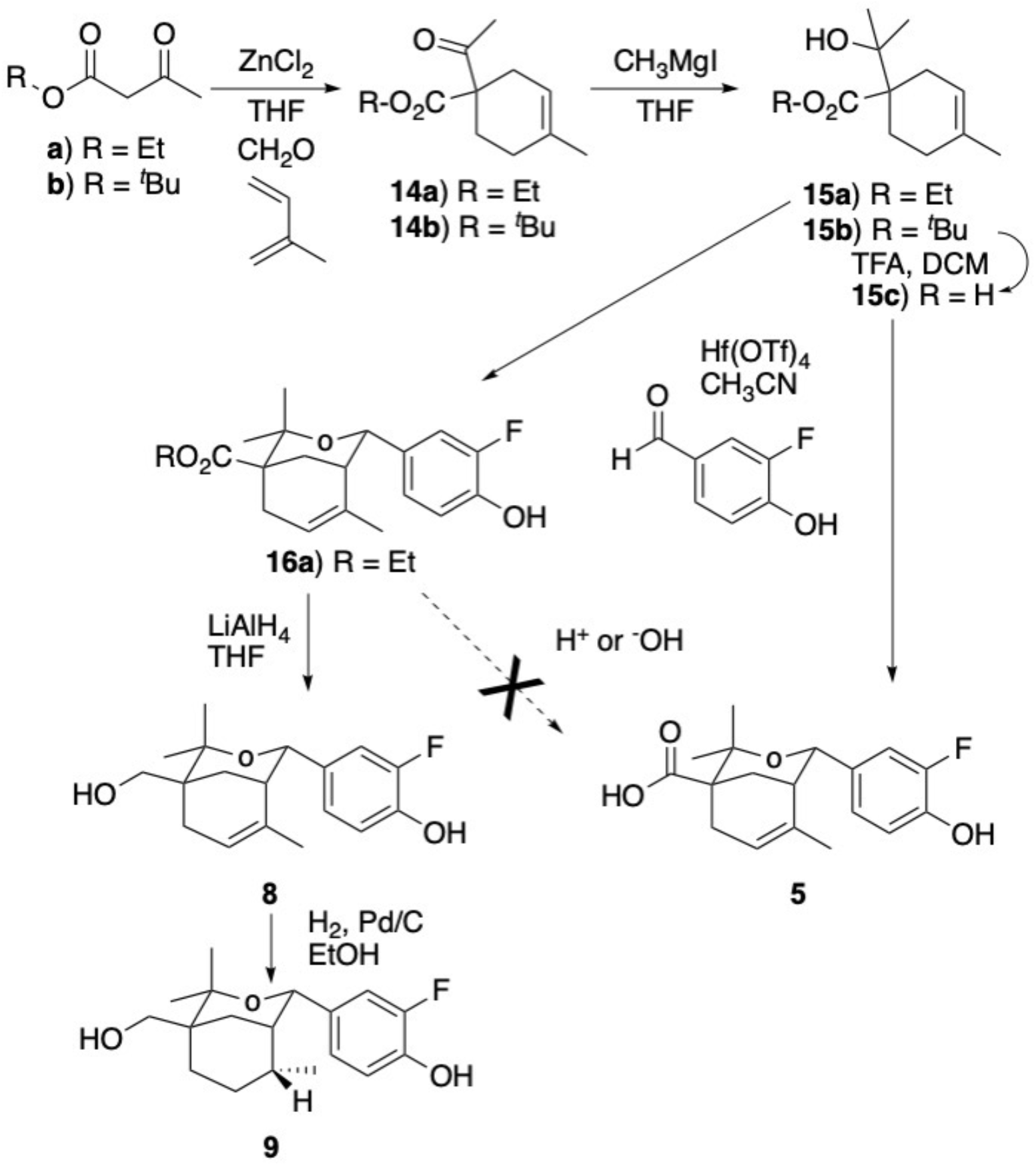
General procedure for the synthesis of compounds **5**, **8** and **9**.

**Scheme 4.**
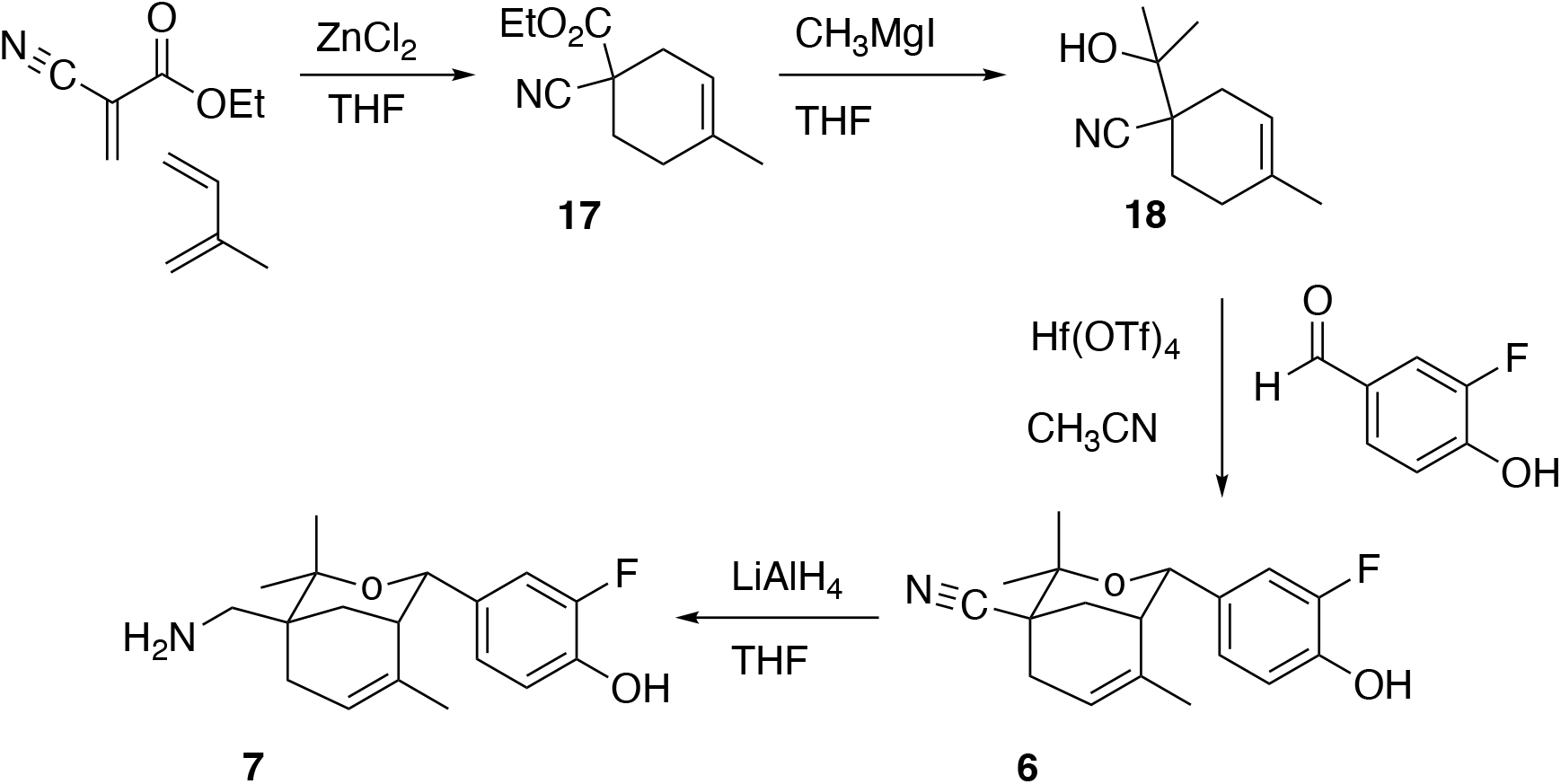
General procedure for the synthesis of compounds **6** and **7**.

## References

1. Mosselman, S., J. Polman, and R. Dijkema, ER beta: identification and characterization of a novel human estrogen receptor. FEBS Lett, 1996. 392(1): p. 49–53 DOI: 10.1016/0014-5793(96)00782-x.

2. Kuiper, G.G., et al., Comparison of the ligand binding specificity and transcript tissue distribution of estrogen receptors alpha and beta. Endocrinology, 1997. 138(3): p. 863–70 DOI: 10.1210/endo.138.3.4979.

3. Hall, J.M. and D.P. McDonnell, The estrogen receptor beta-isoform (ERbeta) of the human estrogen receptor modulates ERalpha transcriptional activity and is a key regulator of the cellular response to estrogens and antiestrogens. Endocrinology, 1999. 140(12): p. 5566–78 DOI: 10.1210/endo.140.12.7179.

4. Koehler, K.F., et al., Reflections on the discovery and significance of estrogen receptor beta. Endocr Rev, 2005. 26(3): p. 465–78 DOI: 10.1210/er.2004-0027.

5. Stauffer, S.R., et al., Pyrazole ligands: structure-affinity/activity relationships and estrogen receptor-alpha-selective agonists. J Med Chem, 2000. 43(26): p. 4934–47 DOI: 10.1021/jm000170m.

6. Meyers, M.J., et al., Estrogen receptor-beta potency-selective ligands: structure-activity relationship studies of diarylpropionitriles and their acetylene and polar analogues. J Med Chem, 2001. 44(24): p. 4230–51 DOI: 10.1021/jm010254a.

7. Fang, H., et al., Structure-activity relationships for a large diverse set of natural, synthetic, and environmental estrogens. Chem Res Toxicol, 2001. 14(3): p. 280–94 DOI: 10.1021/tx000208y.

8. Swaby, R.F., C.G. Sharma, and V.C. Jordan, SERMs for the treatment and prevention of breast cancer. Rev Endocr Metab Disord, 2007. 8(3): p. 229–39 DOI: 10.1007/s11154-007-9034-4.

9. Early Breast Cancer Trialists’ Collaborative, G., Effects of chemotherapy and hormonal therapy for early breast cancer on recurrence and 15-year survival: an overview of the randomised trials. Lancet, 2005. 365(9472): p. 1687–717 DOI: 10.1016/S0140-6736(05)66544-0.

10. Di Leo, A., et al., Final overall survival: fulvestrant 500 mg vs 250 mg in the randomized CONFIRM trial. J Natl Cancer Inst, 2014. 106(1): p. djt337 DOI: 10.1093/jnci/djt337.

11. Andrahennadi, S., et al., Efficacy of Fulvestrant in Women with Hormone-Resistant Metastatic Breast Cancer (mBC): A Canadian Province Experience. Cancers (Basel), 2021. 13(16)DOI: 10.3390/cancers13164163.

12. Jeselsohn, R., et al., ESR1 mutations-a mechanism for acquired endocrine resistance in breast cancer. Nat Rev Clin Oncol, 2015. 12(10): p. 573–83 DOI: 10.1038/nrclinonc.2015.117.

13. Rasha, F., M. Sharma, and K. Pruitt, Mechanisms of endocrine therapy resistance in breast cancer. Mol Cell Endocrinol, 2021. 532: p. 111322 DOI: 10.1016/j.mce.2021.111322.

14. Pepermans, R.A. and E.R. Prossnitz, ERalpha-targeted endocrine therapy, resistance and the role of GPER. Steroids, 2019. 152: p. 108493 DOI: 10.1016/j.steroids.2019.108493.

15. Patel, R., et al., An emerging generation of endocrine therapies in breast cancer: a clinical perspective. NPJ Breast Cancer, 2023. 9(1): p. 20 DOI: 10.1038/s41523-023-00523-4.

16. Haines, C.N., S.E. Wardell, and D.P. McDonnell, Current and emerging estrogen receptor-targeted therapies for the treatment of breast cancer. Essays Biochem, 2021. 65(6): p. 985–1001 DOI: 10.1042/EBC20200174.

17. Campone, M., et al., Vepdegestrant, a PROTAC Estrogen Receptor Degrader, in Advanced Breast Cancer. N Engl J Med, 2025. 393(6): p. 556–568 DOI: 10.1056/NEJMoa2505725.

18. Sibley, R., et al., A novel estrogen receptor ligand template. Bioorg Med Chem Lett, 2003. 13(11): p. 1919–22 DOI: 10.1016/s0960-894x(03)00307-x.

19. Hamann, L.G., et al., Structure-activity relationships and sub-type selectivity in an oxabicyclic estrogen receptor alpha/beta agonist scaffold. Bioorg Med Chem Lett, 2005. 15(5): p. 1463–6 DOI: 10.1016/j.bmcl.2004.12.077.

20. Hsieh, R.W., et al., Identification of ligands with bicyclic scaffolds provides insights into mechanisms of estrogen receptor subtype selectivity. J Biol Chem, 2006. 281(26): p. 17909–19 DOI: 10.1074/jbc.M513684200.

21. Revankar, C.M., et al., A Selective Ligand for Estrogen Receptor Proteins Discriminates Rapid and Genomic Signaling. Cell Chem Biol, 2019. 26(12): p. 1692–1702 e5 DOI: 10.1016/j.chembiol.2019.10.009.

22. Brzozowski, A.M., et al., Molecular basis of agonism and antagonism in the oestrogen receptor. Nature, 1997. 389(6652): p. 753–8 DOI: 10.1038/39645.

23. Liu, J., et al., Fulvestrant-3 Boronic Acid (ZB716): An Orally Bioavailable Selective Estrogen Receptor Downregulator (SERD). J Med Chem, 2016. 59(17): p. 8134–40 DOI: 10.1021/acs.jmedchem.6b00753.

24. Katzenellenbogen, J.A., B.W. O’Malley, and B.S. Katzenellenbogen, Tripartite steroid hormone receptor pharmacology: interaction with multiple effector sites as a basis for the cell- and promoter-specific action of these hormones. Mol Endocrinol, 1996. 10(2): p. 119–31 DOI: 10.1210/mend.10.2.8825552.

25. Heldring, N., et al., Estrogen receptors: how do they signal and what are their targets. Physiol Rev, 2007. 87(3): p. 905–31 DOI: 10.1152/physrev.00026.2006.

26. Yamaguchi, Y., et al., Tumor-stromal interaction through the estrogen-signaling pathway in human breast cancer. Cancer Res, 2005. 65(11): p. 4653–62 DOI: 10.1158/0008-5472.CAN-04-3236.

27. Gough, S.M., et al., Oral Estrogen Receptor PROTAC Vepdegestrant (ARV-471) Is Highly Efficacious as Monotherapy and in Combination with CDK4/6 or PI3K/mTOR Pathway Inhibitors in Preclinical ER+ Breast Cancer Models. Clin Cancer Res, 2024. 30(16): p. 3549–3563 DOI: 10.1158/1078-0432.CCR-23-3465.

28. Hamilton, E.P., et al., PROteolysis TArgeting Chimera (PROTAC) Estrogen Receptor Degraders for Treatment of Estrogen Receptor-Positive Advanced Breast Cancer. Target Oncol, 2025. 20(3): p. 431–444 DOI: 10.1007/s11523-025-01137-5.

29. Lu, A.S., et al., Proteolytic Targeting Chimeras with Specificity for Plasma Membrane and Intracellular Estrogen Receptors. Mol Pharm, 2021. 18(3): p. 1455–1469 DOI: 10.1021/acs.molpharmaceut.1c00018.

30. Chu, K.C., H.J. Liu, and J.L. Zhu, A New and Efficient Total Synthesis of (±)-Laurencenone C. Synlett, 2010(20): p. 3061–3064 DOI: 10.1055/s-0030-1259052.

31. Saxon, T.A., Synthesis and optimization of novel oxabicyclic scaffolds for selective modulation of estrogen receptor targets., in Chemistry. 2020, New Mexico State University. p. 161.

