## Supplemental Data for "Synthesis of substituted oxabicyclic phenol derivatives with increased affinity and activity for the classical estrogen receptor ERα"

Abbreviations: C_6_F_6_, hexafluorobenzene; CDCl_3_, deuterochloroform; CD_3_OD, methanol‑d_4_

**Table of Contents**

**Figure S1.** ^1^H‑NMR, ^13^C‑NMR and ^19^F‑NMR spectra for **2**	**3**−**4**

**Figure S2.** ^1^H‑NMR, ^13^C‑NMR and ^19^F‑NMR spectra for **3**	**5**−**6**

**Figure S3.** ^1^H‑NMR and ^13^C‑NMR spectra for **4**	**7**

**Figure S4.** ^1^H‑NMR, ^13^C‑NMR and ^19^F‑NMR spectra for **5**	**8**−**9**

**Figure S5.** ^1^H‑NMR, ^13^C‑NMR and ^19^F‑NMR spectra for **6**	**10**−**11**

**Figure S6.** ^1^H‑NMR, ^13^C‑NMR and ^19^F‑NMR spectra for **7**	**12**−**13**

**Figure S7.** ^1^H‑NMR, ^13^C‑NMR and ^19^F‑NMR spectra for **8**	**14**−**15**

**Figure S8.** ^1^H‑NMR, ^13^C‑NMR and ^19^F‑NMR spectra for **9**	**16**−**17**

**Table S1.** Summary of multi‑step syntheses of compounds **2**‑**9**	**18**

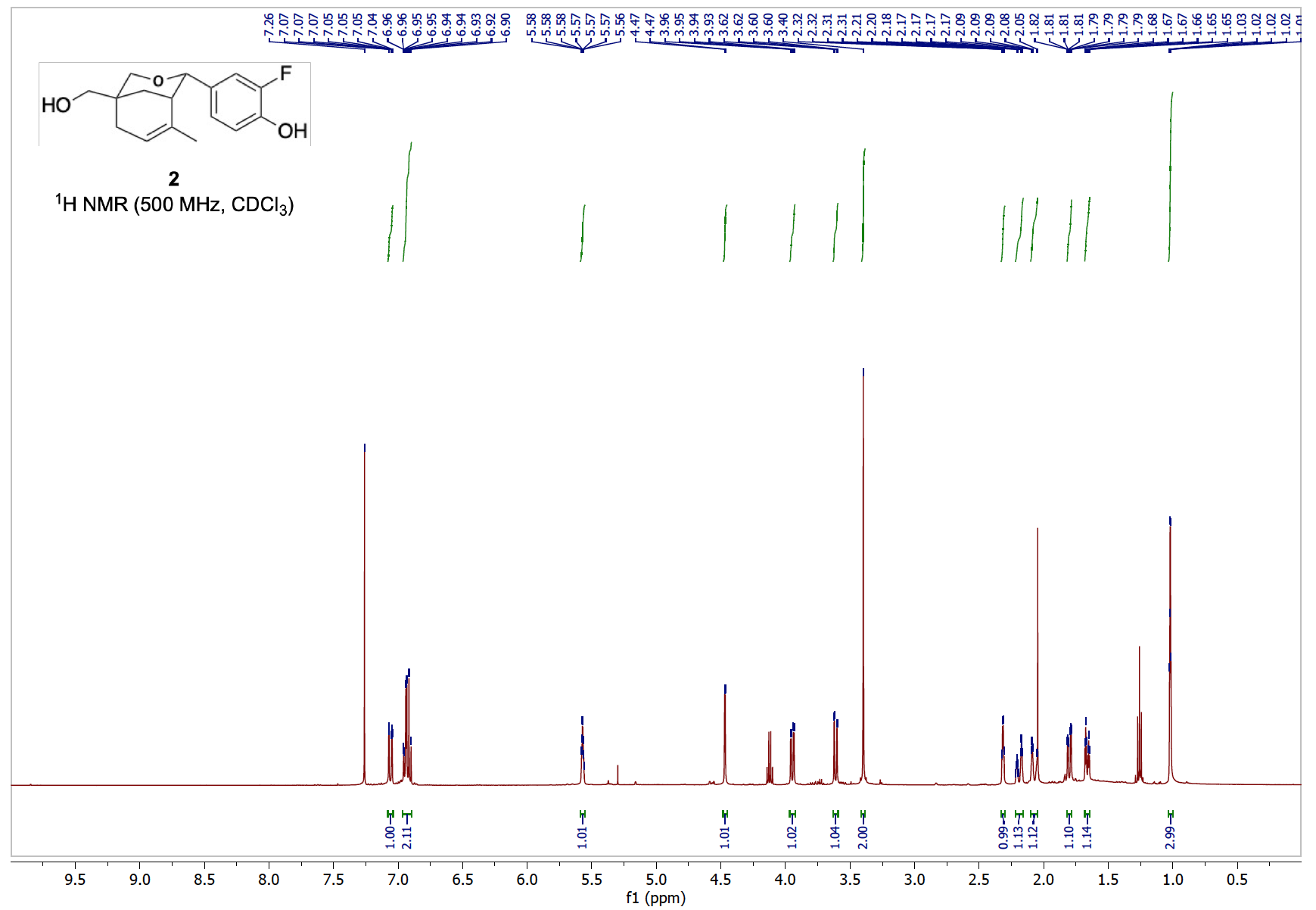

CDCl_3_

**A**

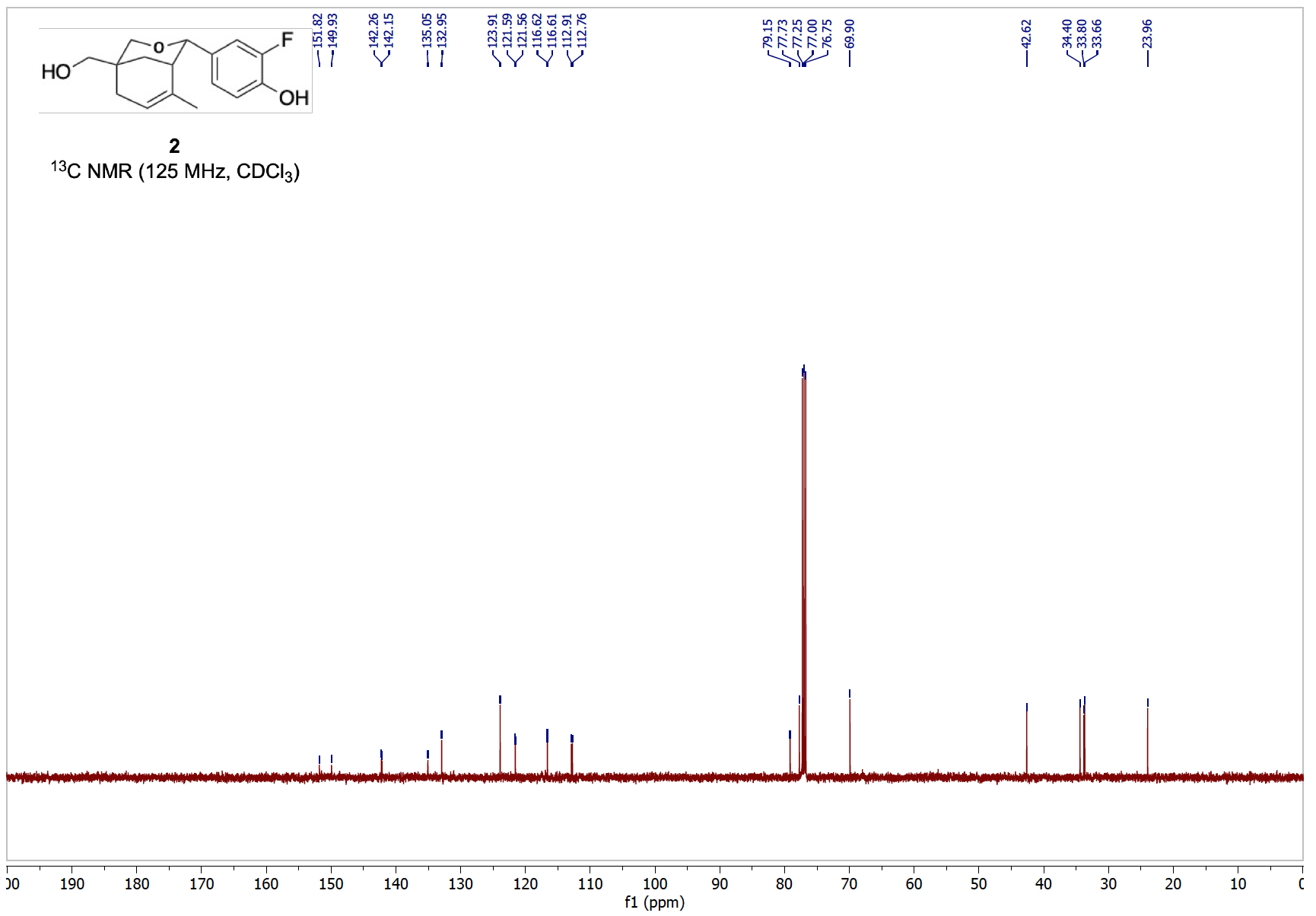

**B**

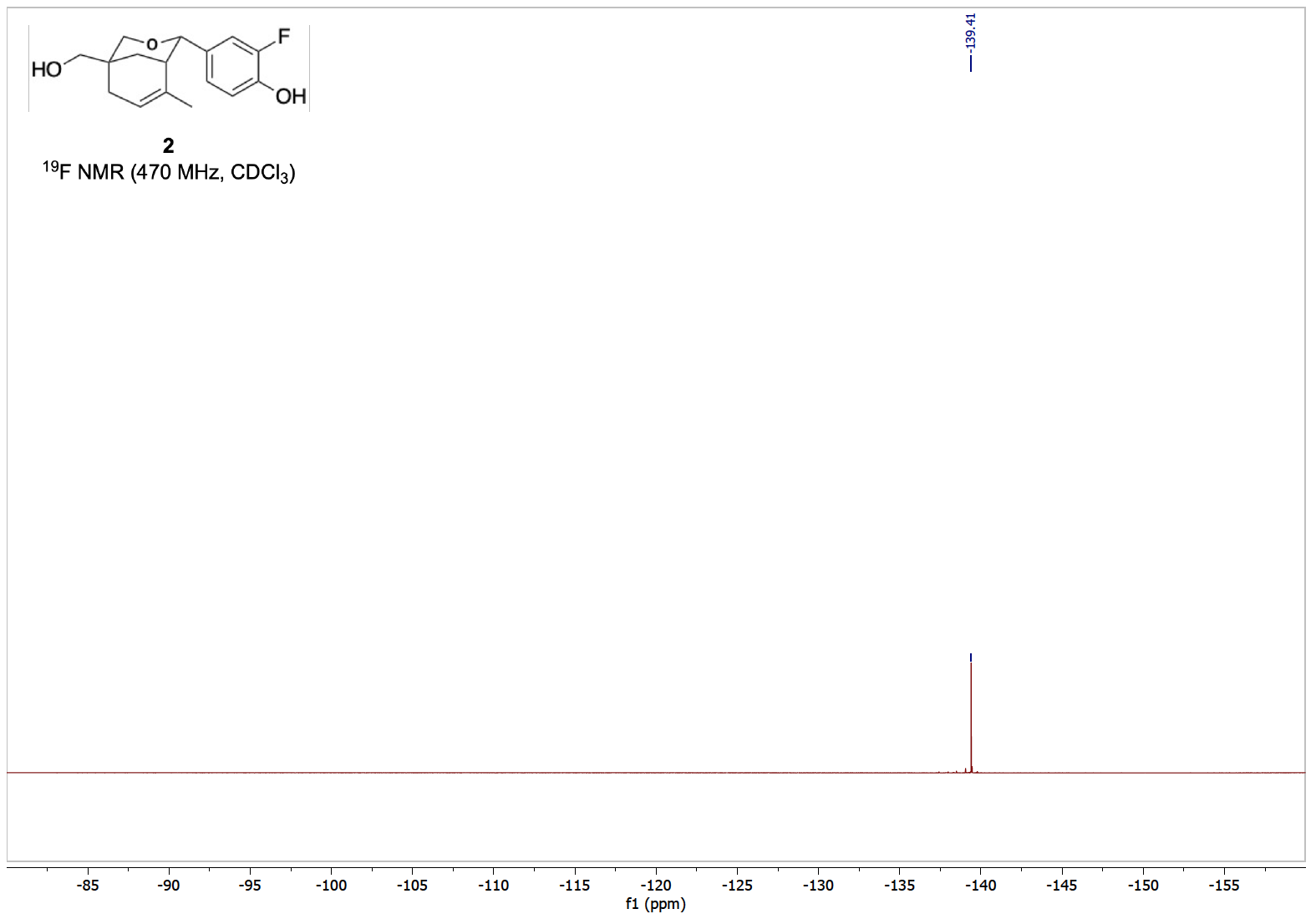

**C**

**Figure S1.** **(A)** ^1^H‑NMR (500 MHz), **(B)** ^13^C‑NMR (125 MHz) and **(C)** ^19^F‑NMR (470 MHz) of compound **2**.

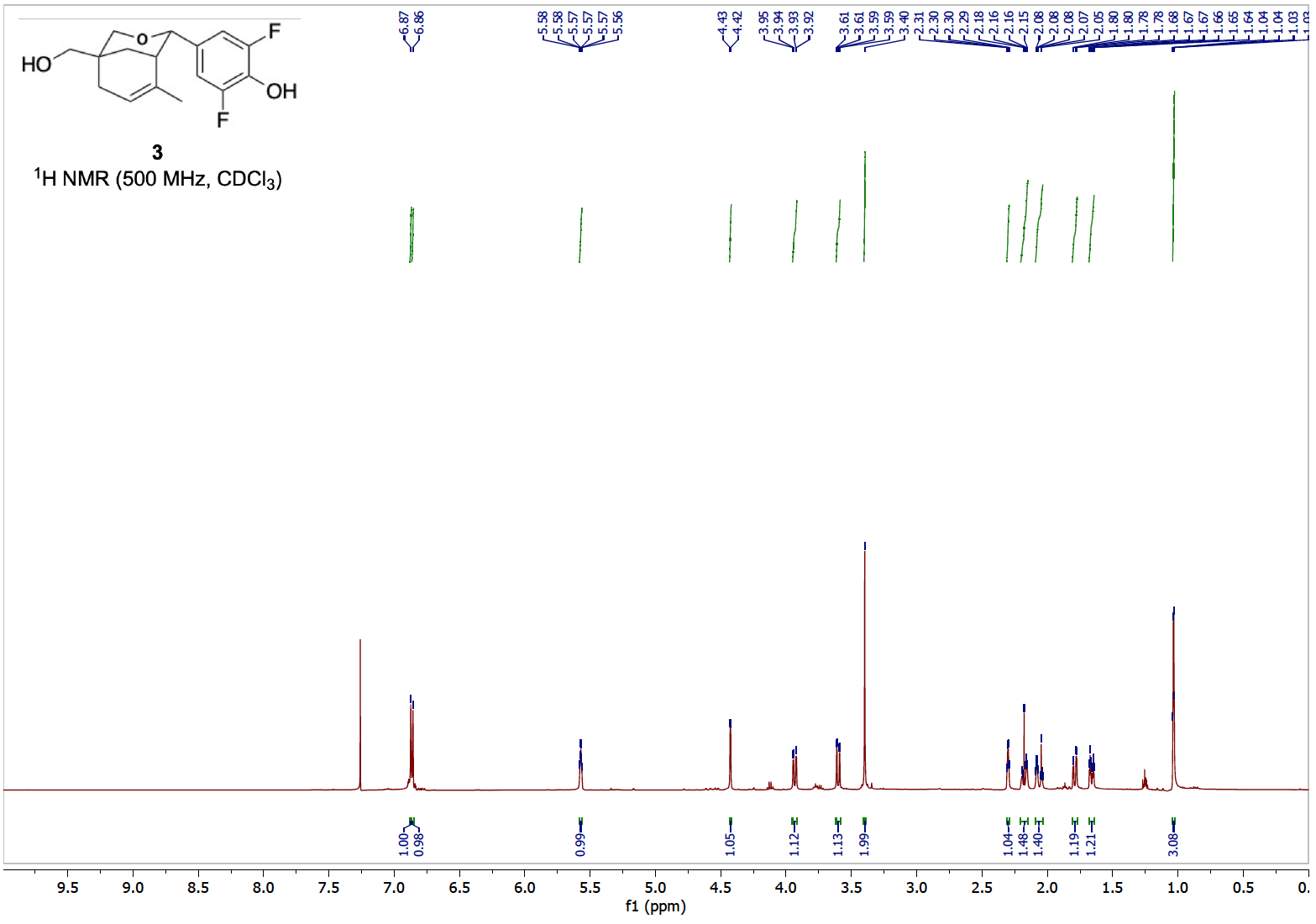

**A**

CDCl_3_

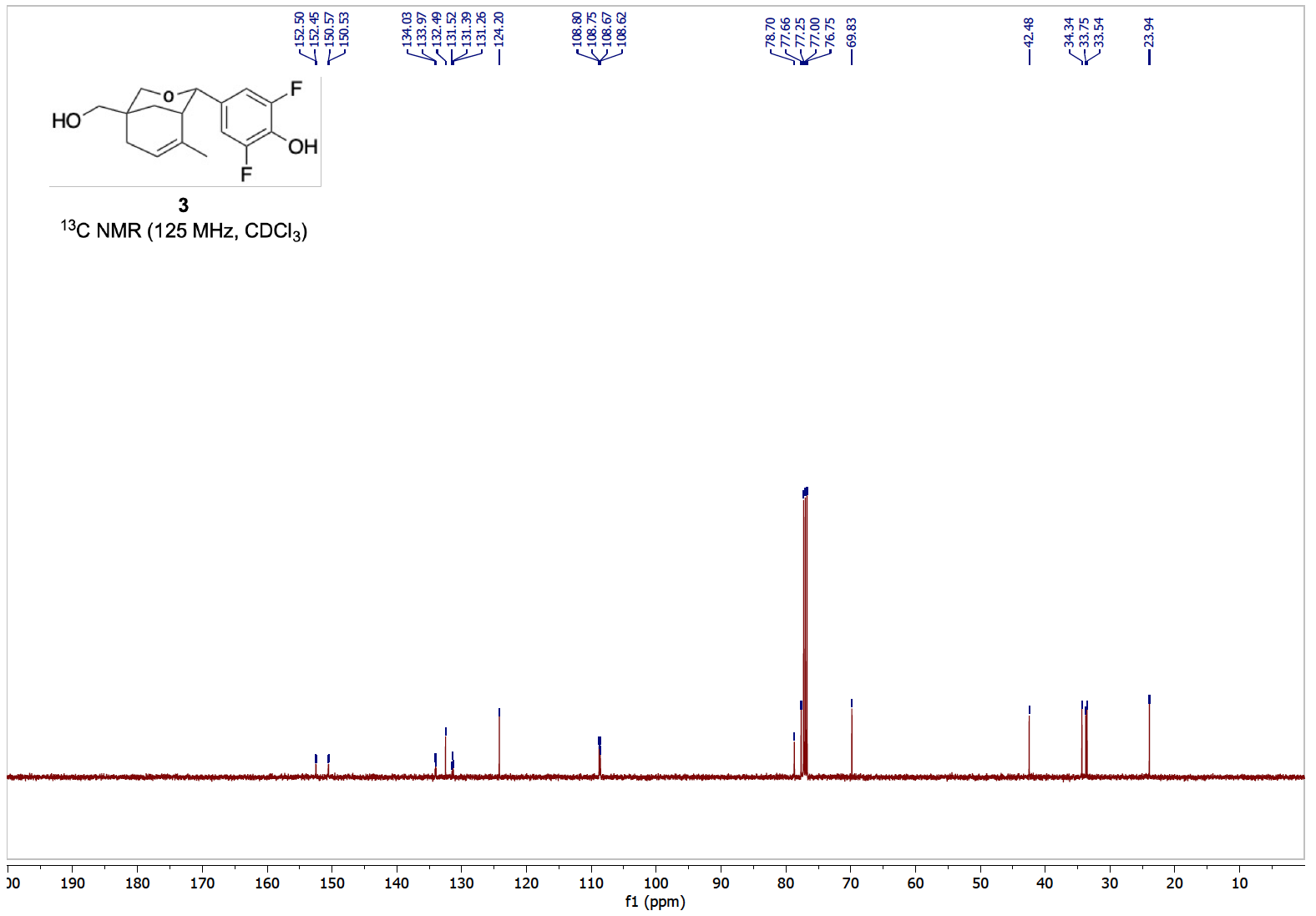

**B**

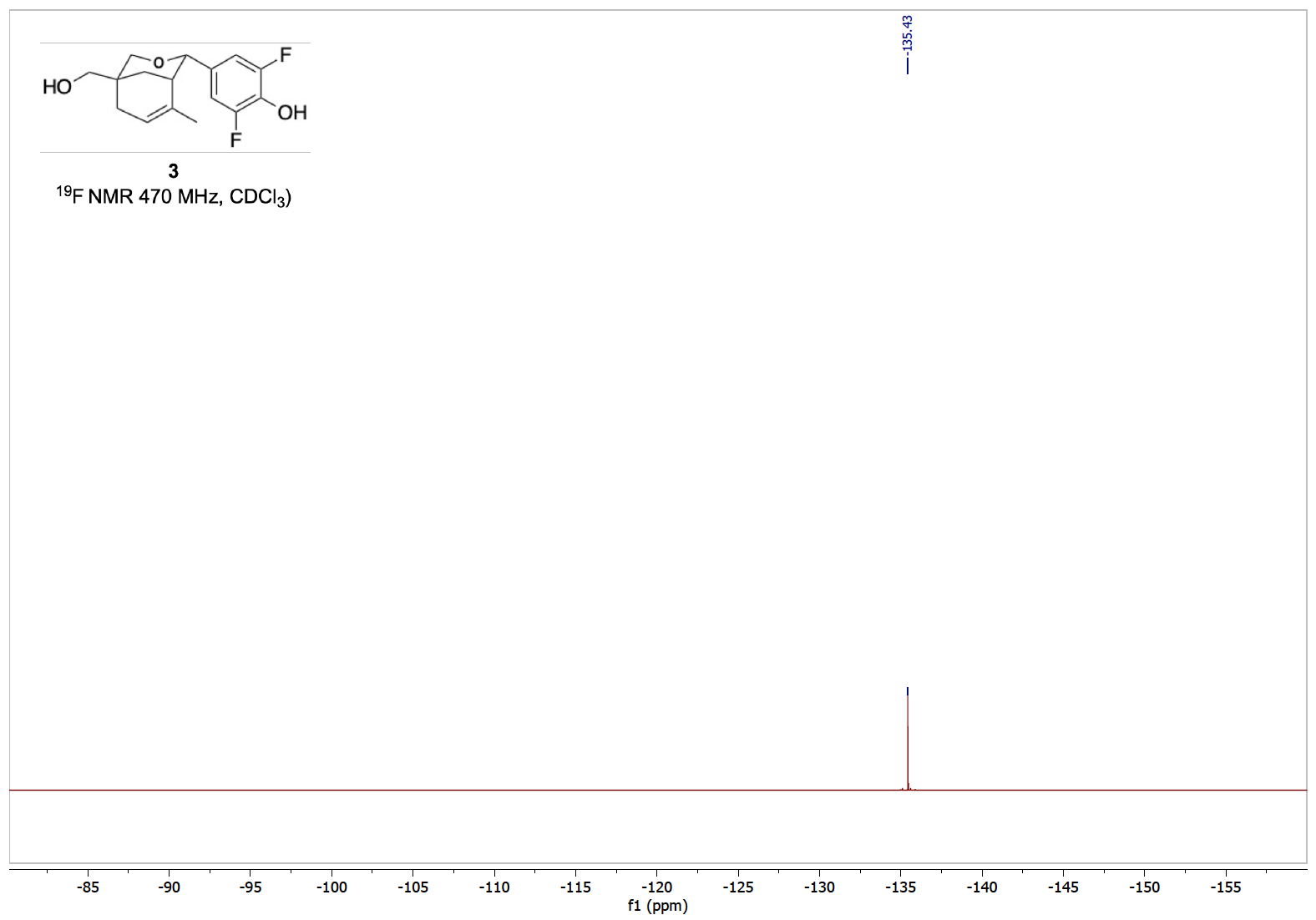

**CC**

**Figure S2.** **(A)** ^1^H‑NMR (500 MHz), **(B)** ^13^C‑NMR (125 MHz) and **(C)** ^19^F‑NMR (470 MHz) of compound **3**.

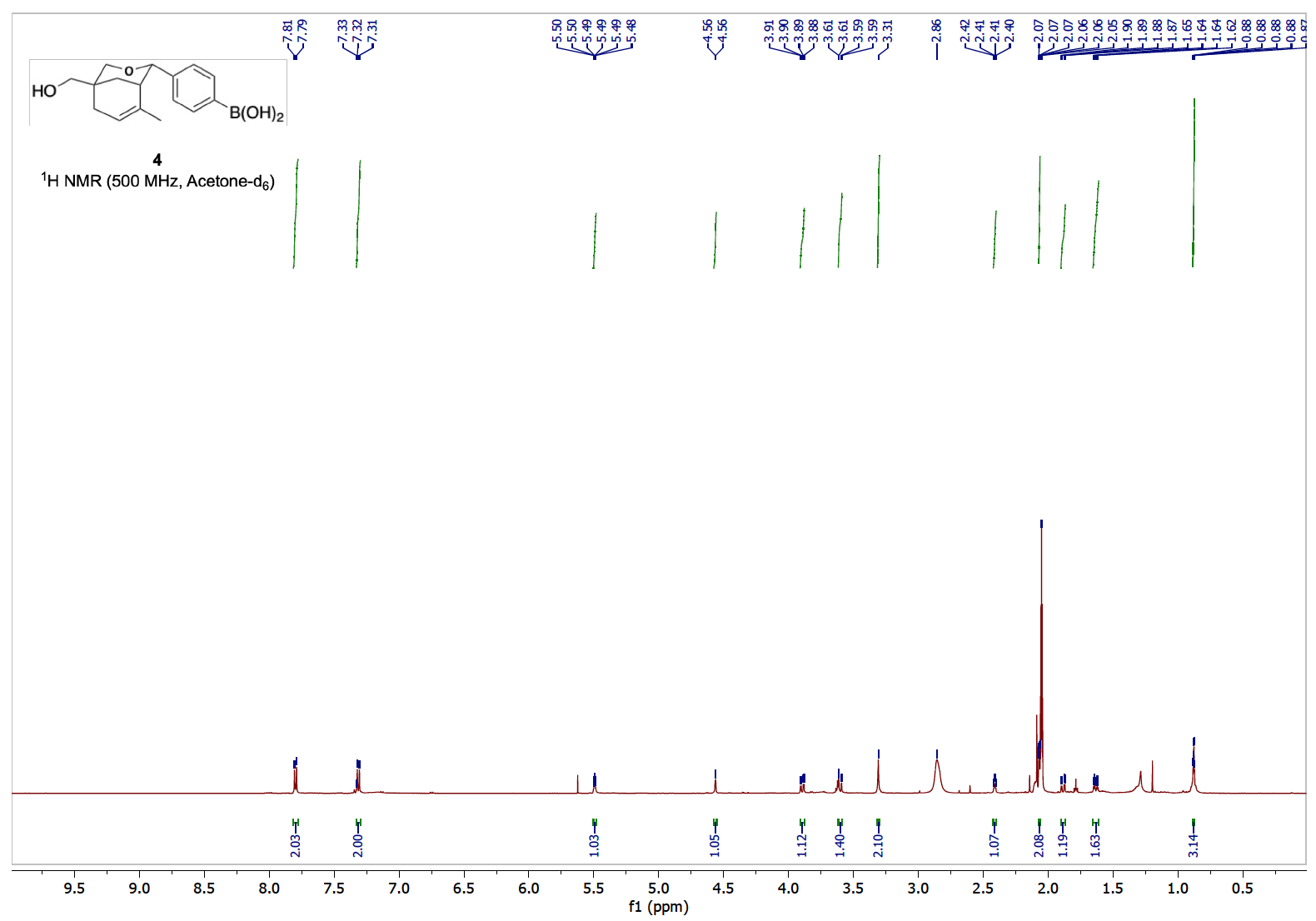

**A**

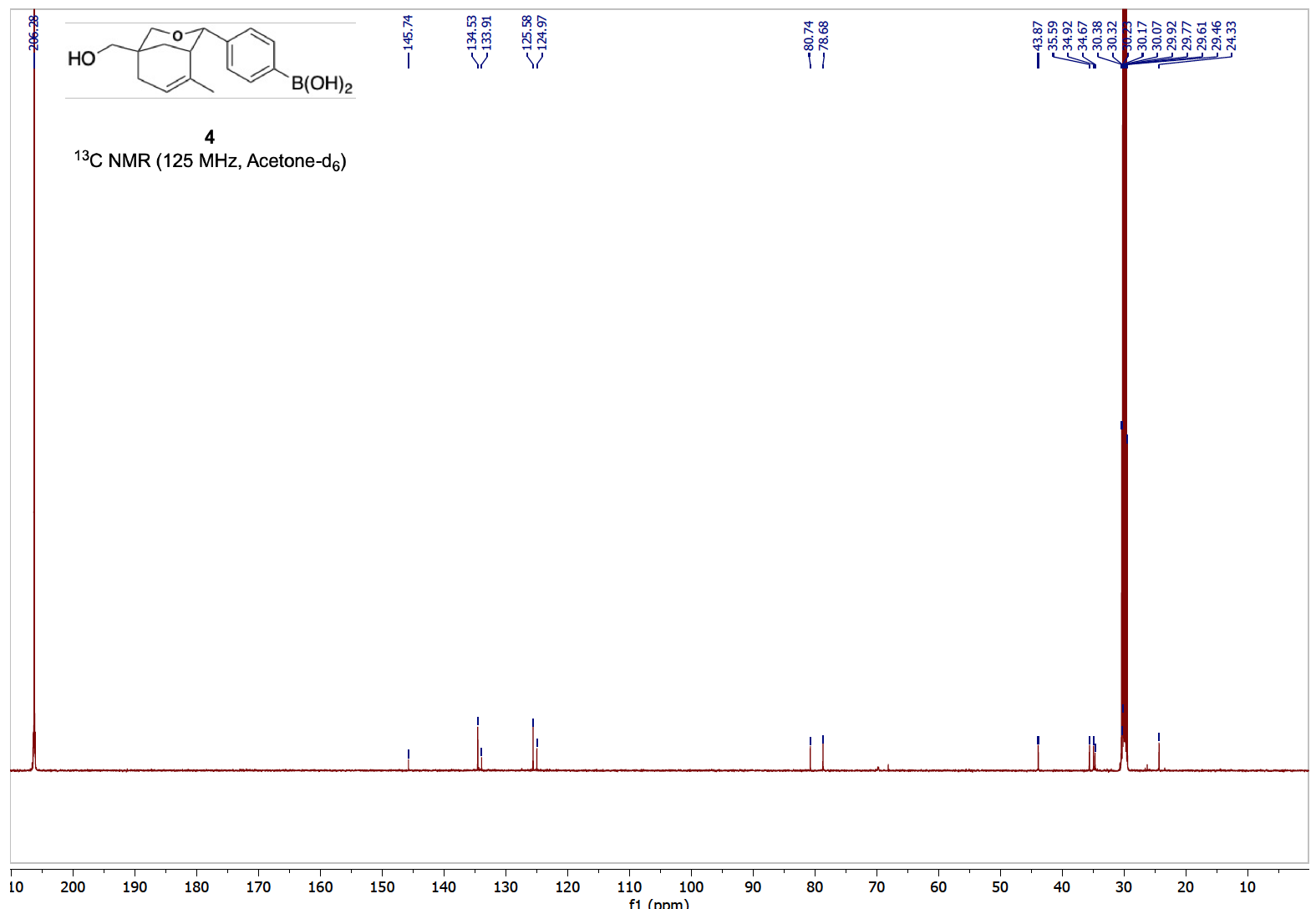

**B**

Acetone‑d_6_

(singlet)

**Figure S3.** **(A)** ^1^H‑NMR (500 MHz) and **(B)** ^13^C‑NMR (125 MHz) of compound **4**.

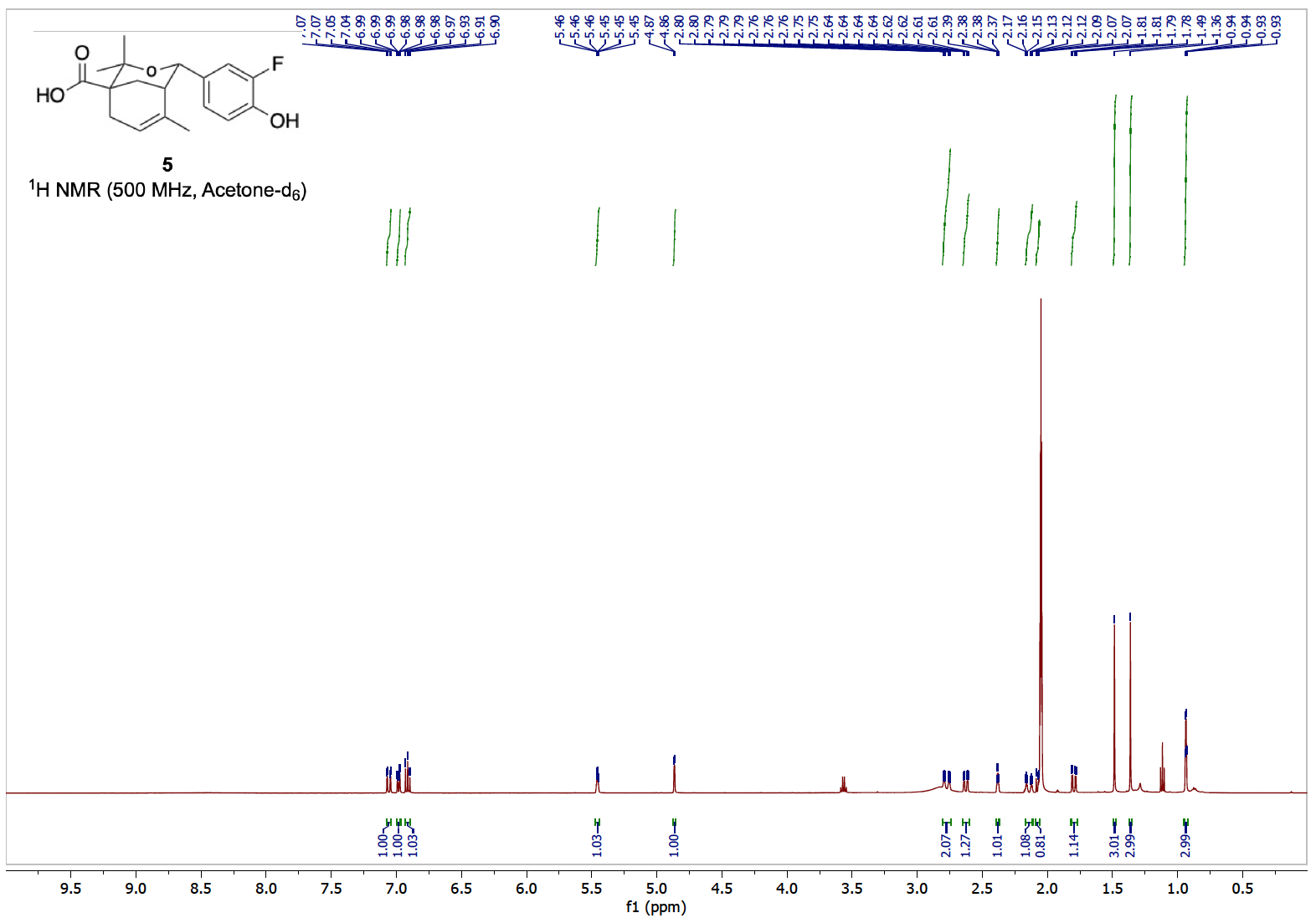

**A**

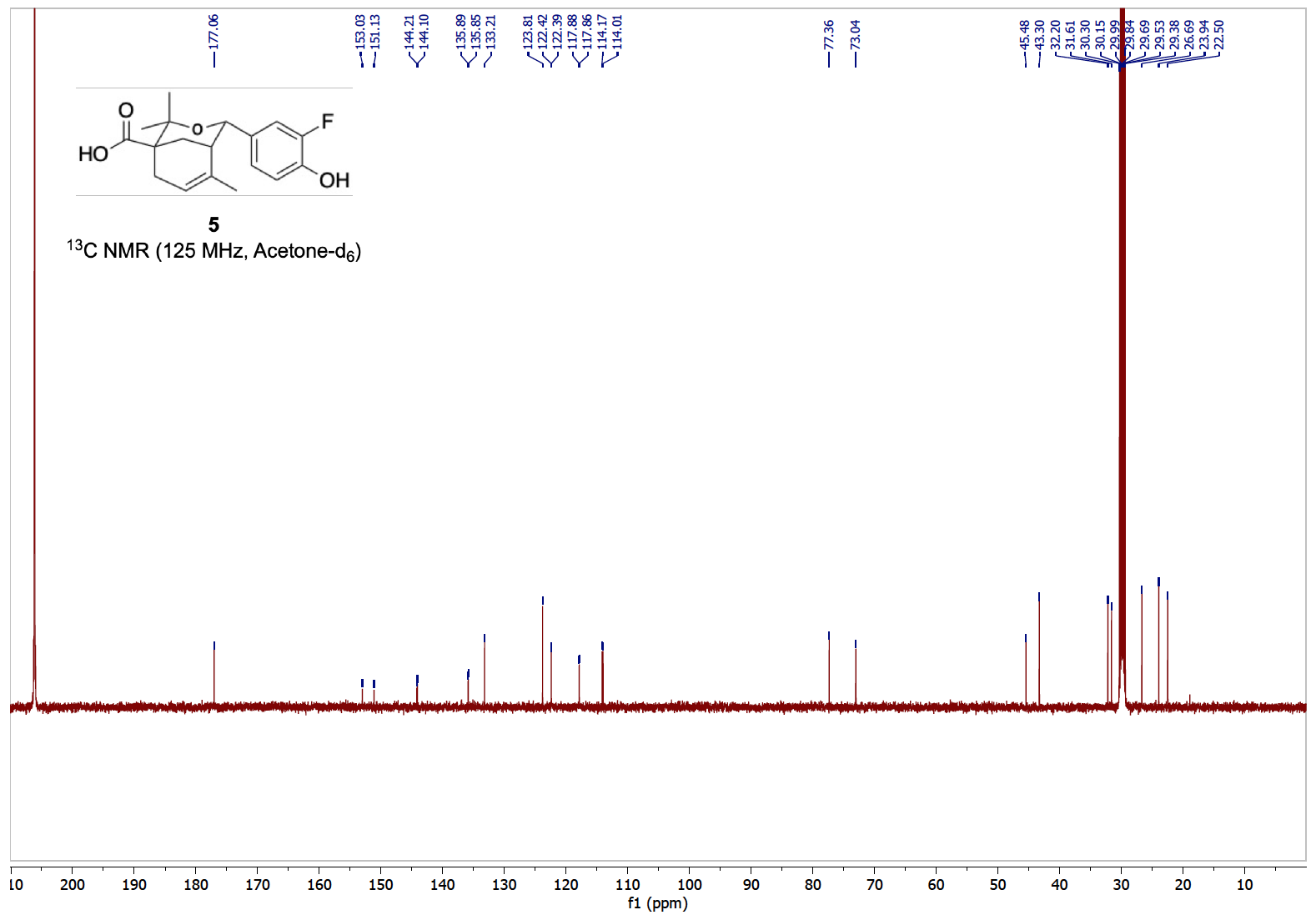

Acetone‑d_6_

(septet)

Acetone‑d_6_

(singlet)

**B**

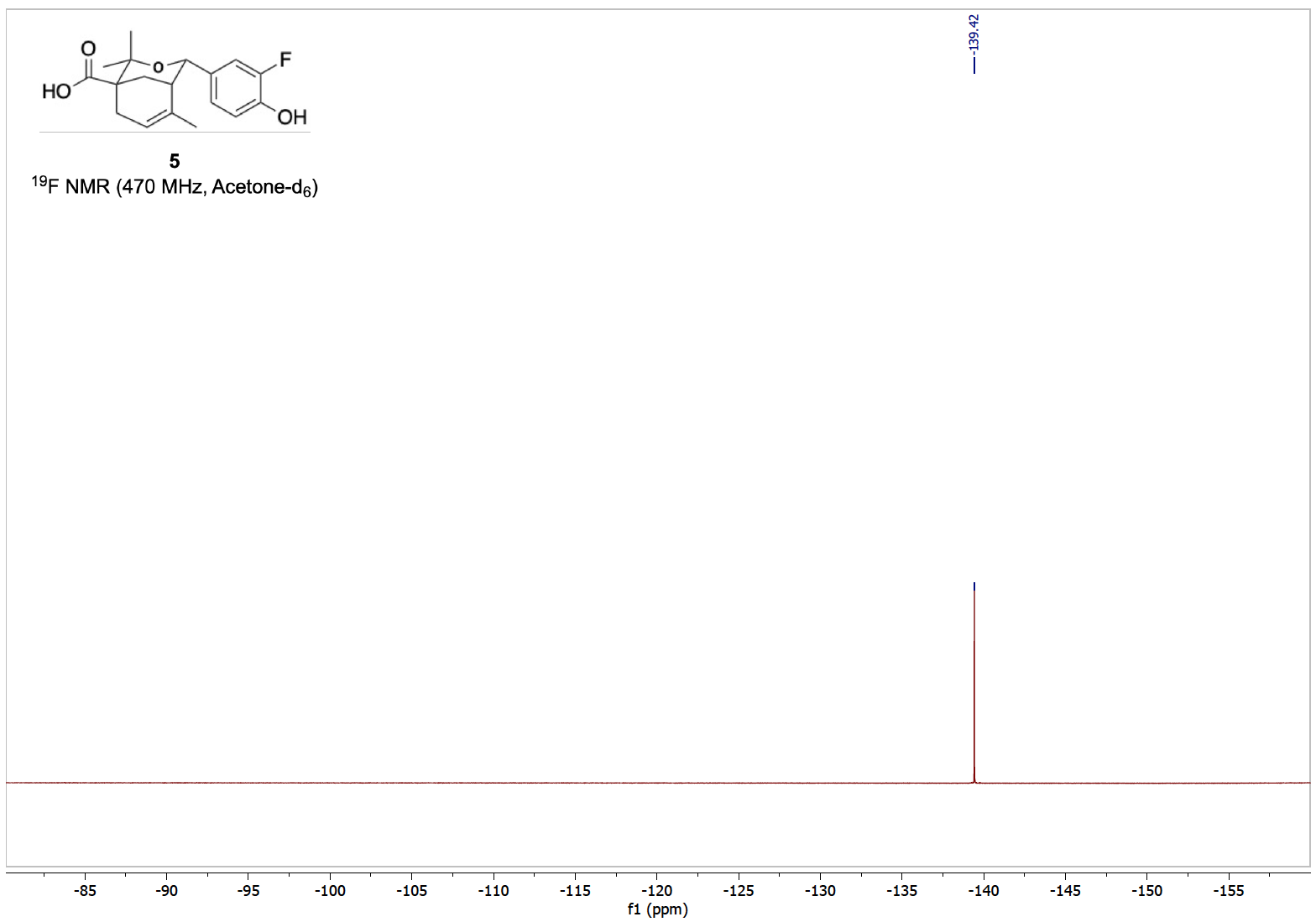

**C**

**Figure S4.** **(A)** ^1^H‑NMR (500 MHz), **(B)** ^13^C‑NMR (125 MHz) and **(C)** ^19^F‑NMR (470 MHz) of compound **5**.

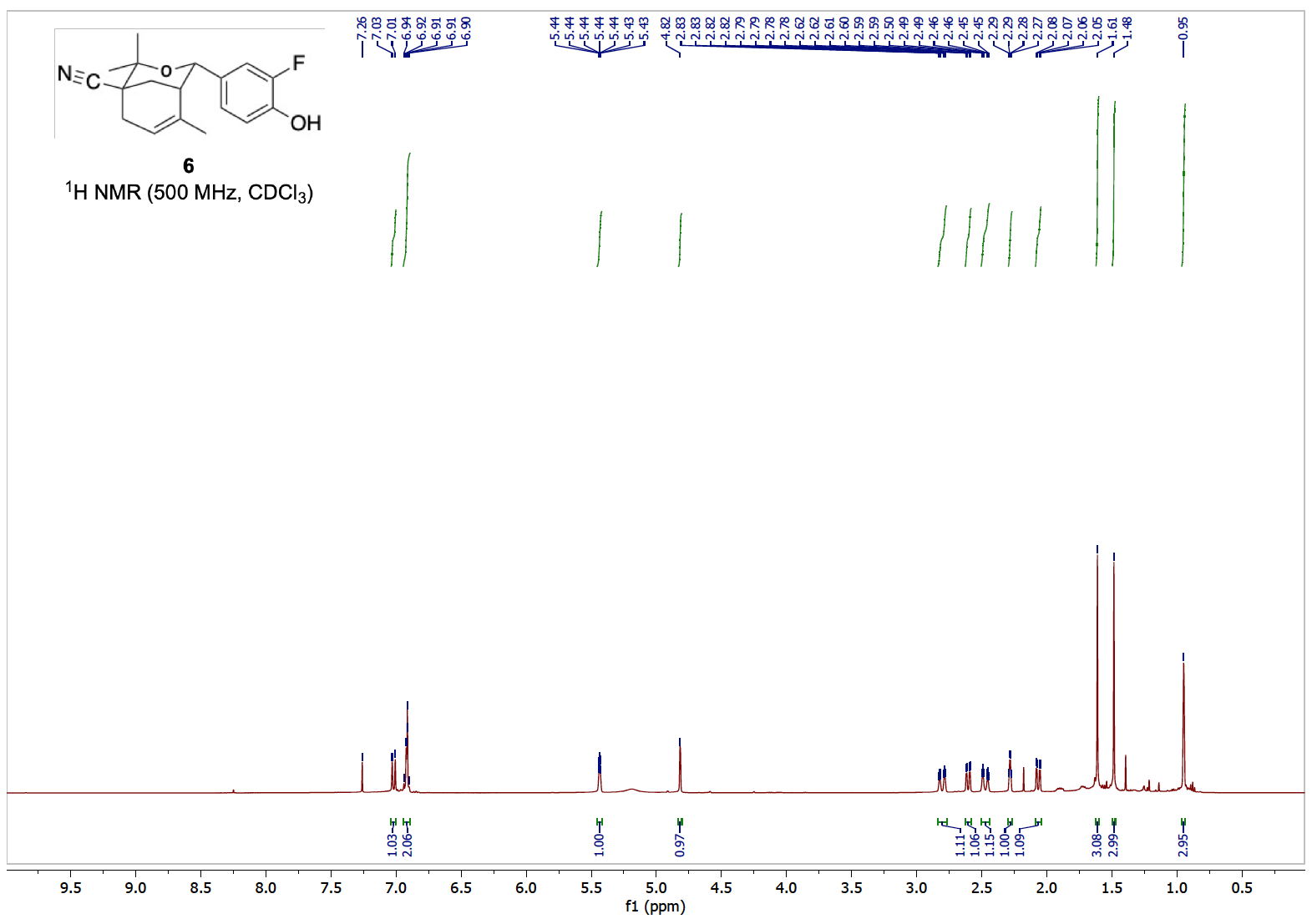

CDCl_3_

**A**

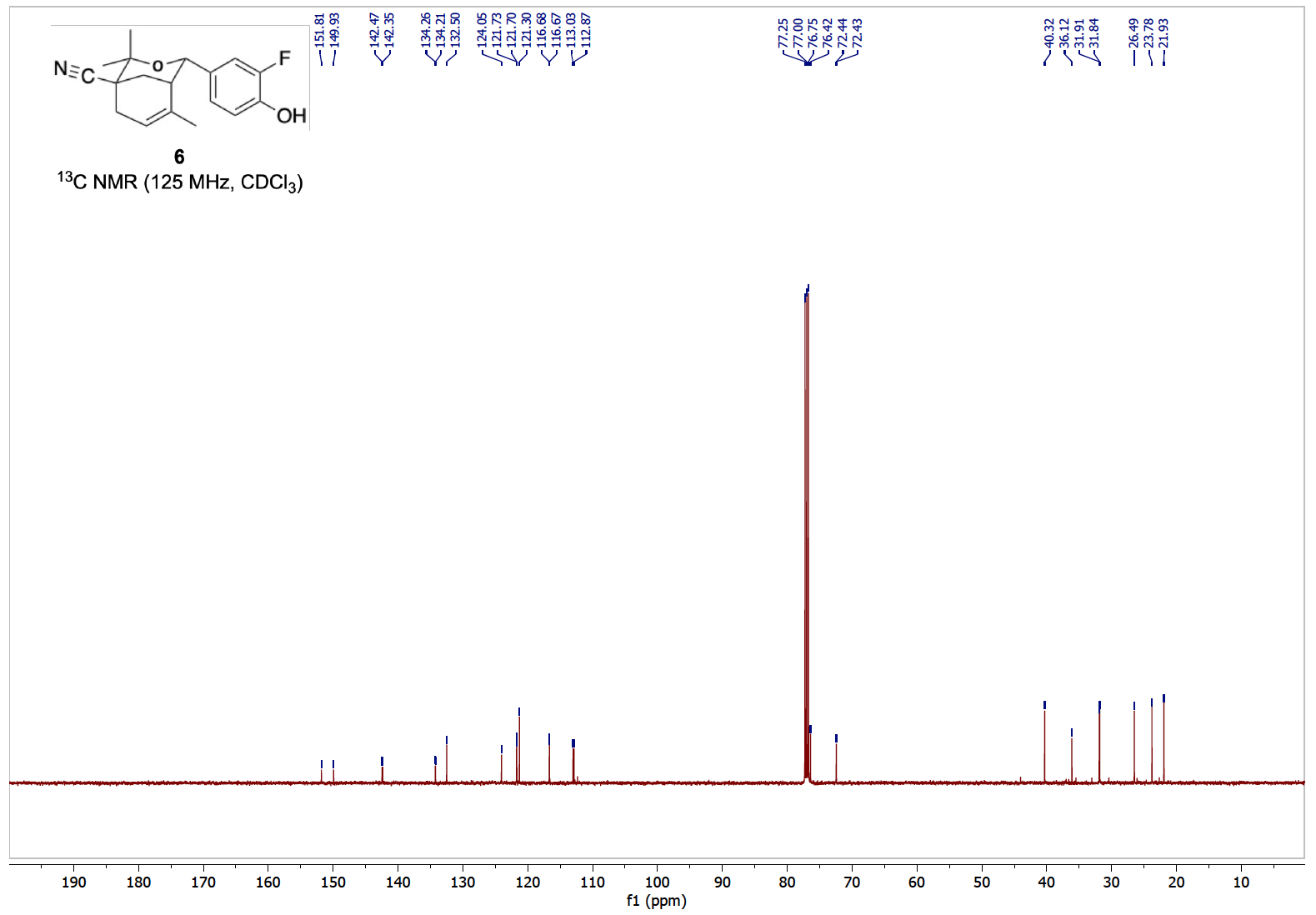

**B**

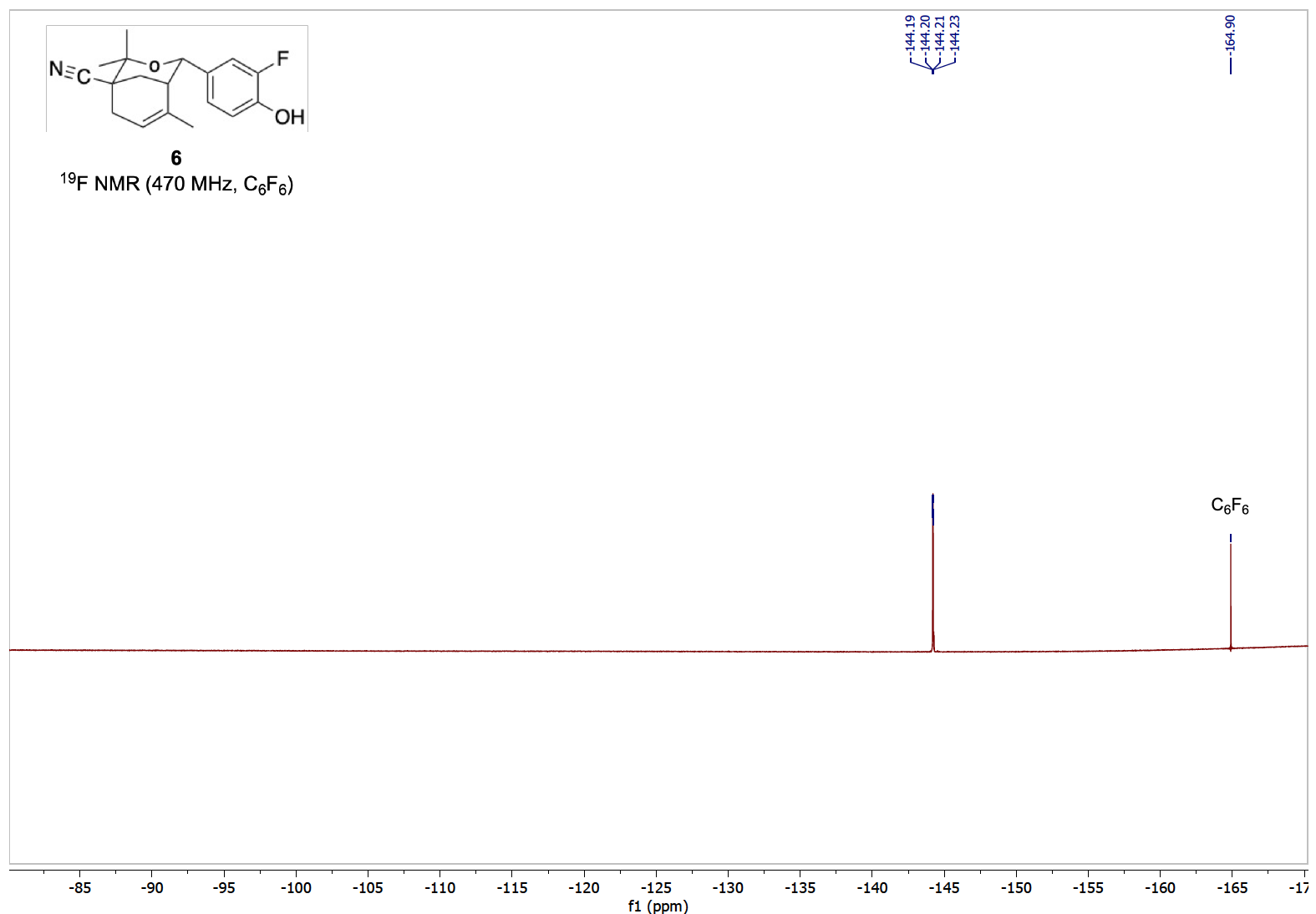

**C**

**Figure S5.** **(A)** ^1^H‑NMR (500 MHz), **(B)** ^13^C‑NMR (125 MHz) and **(C)** ^19^F‑NMR (470 MHz) of compound **6**.

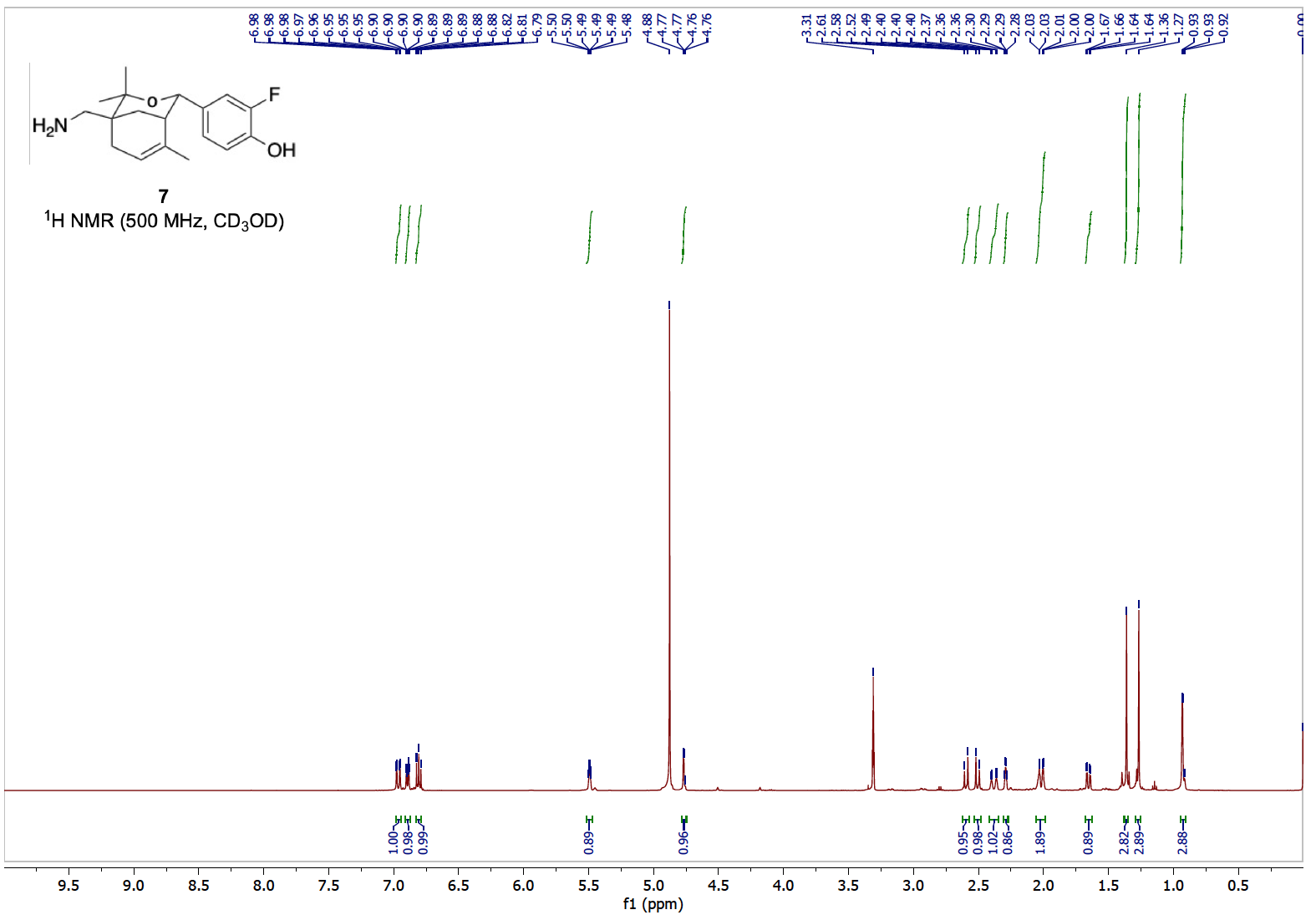

CD_3_OD

**A**

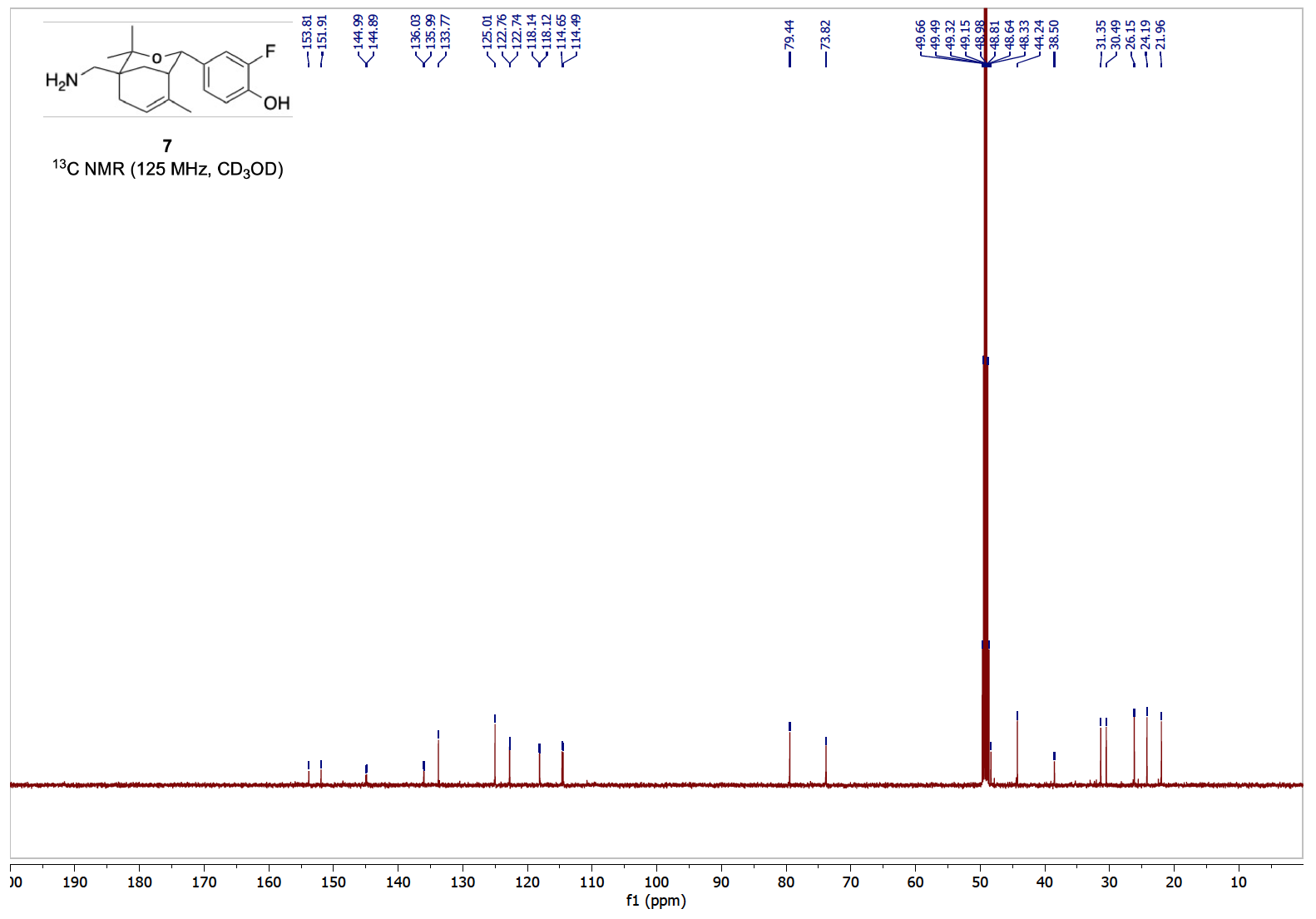

**B**

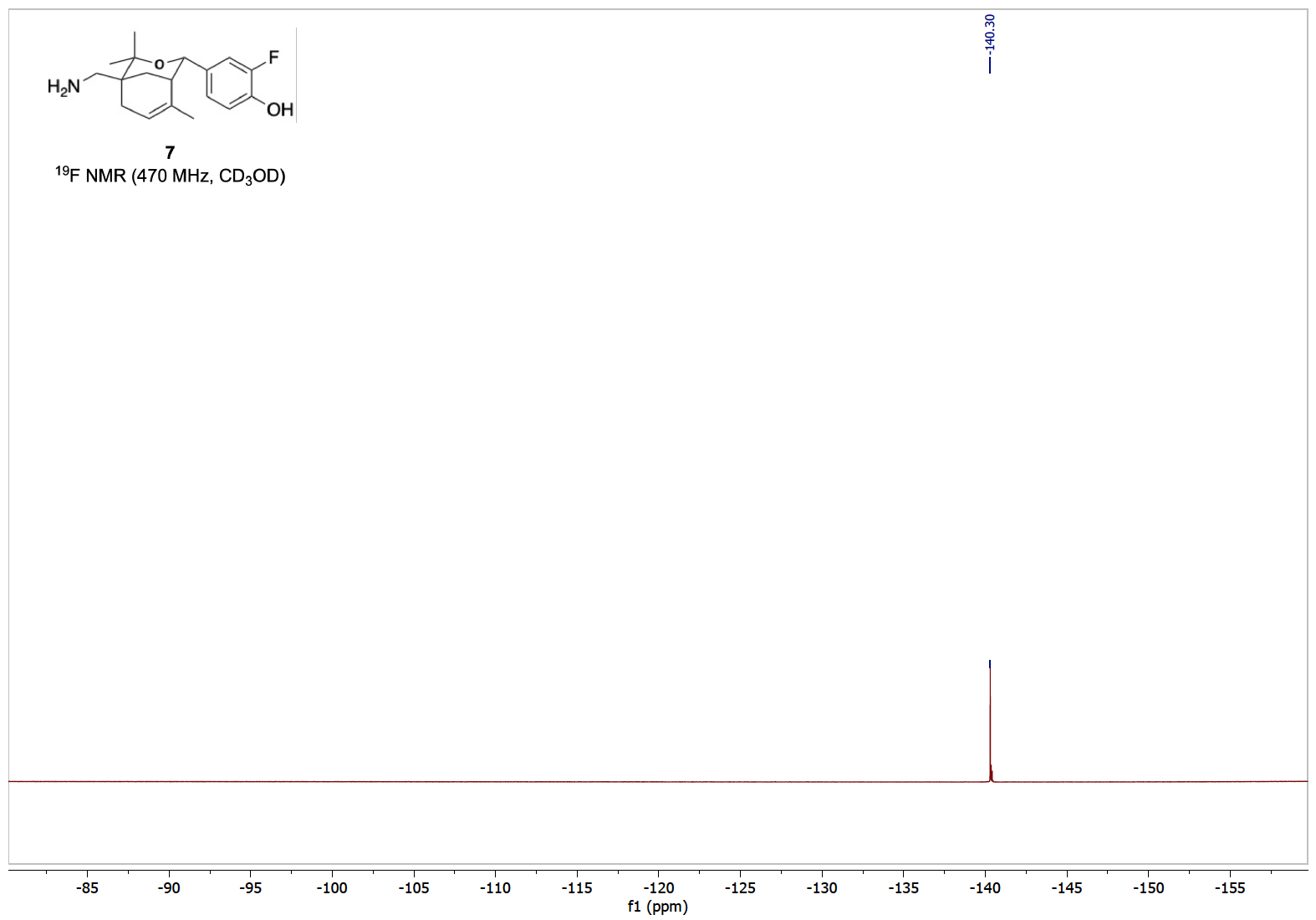

**C**

**Figure S6.** **(A)** ^1^H‑NMR (500 MHz), **(B)** ^13^C‑NMR (125 MHz) and **(C)** ^19^F‑NMR (470 MHz) of compound **7**.

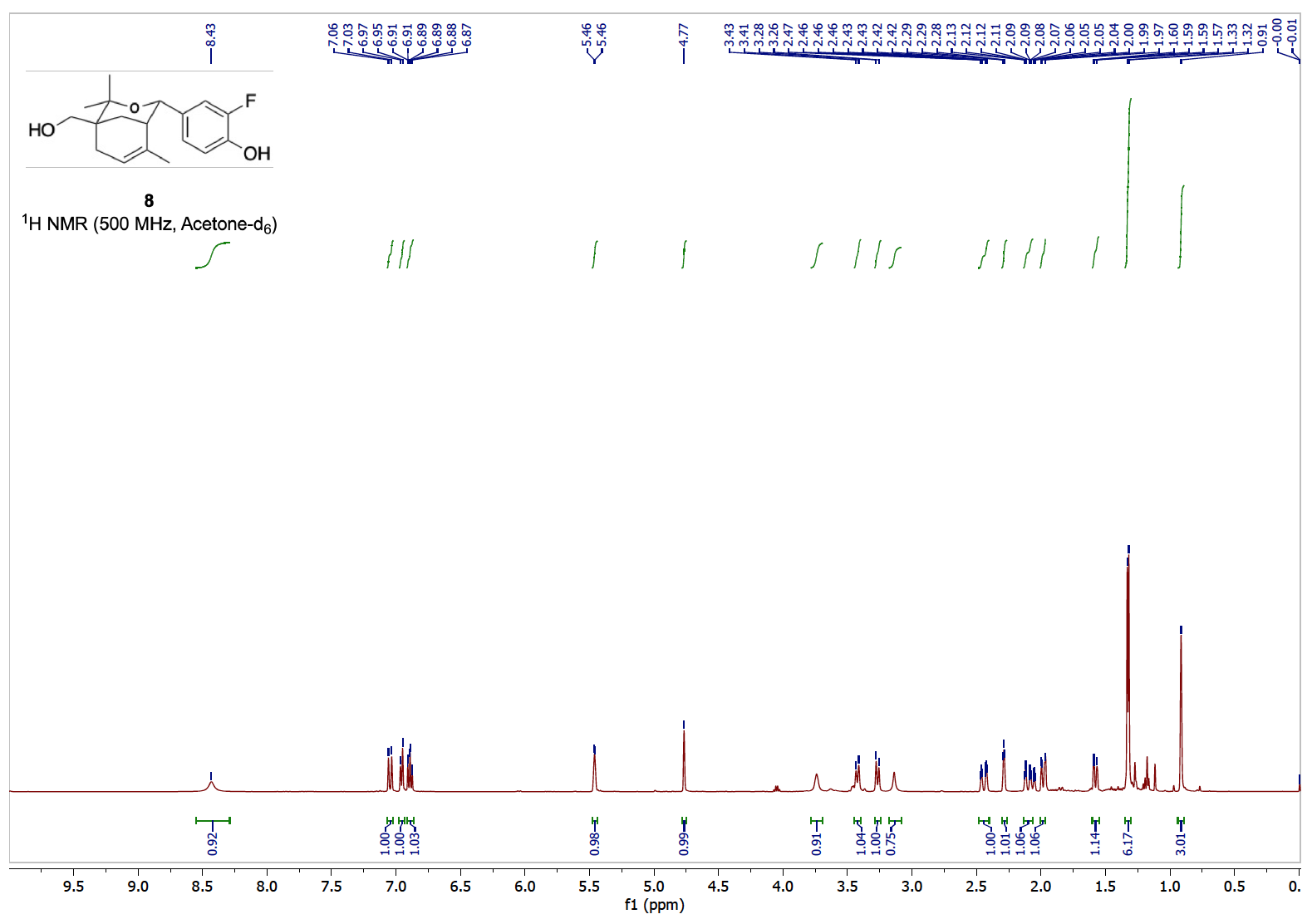

**A**

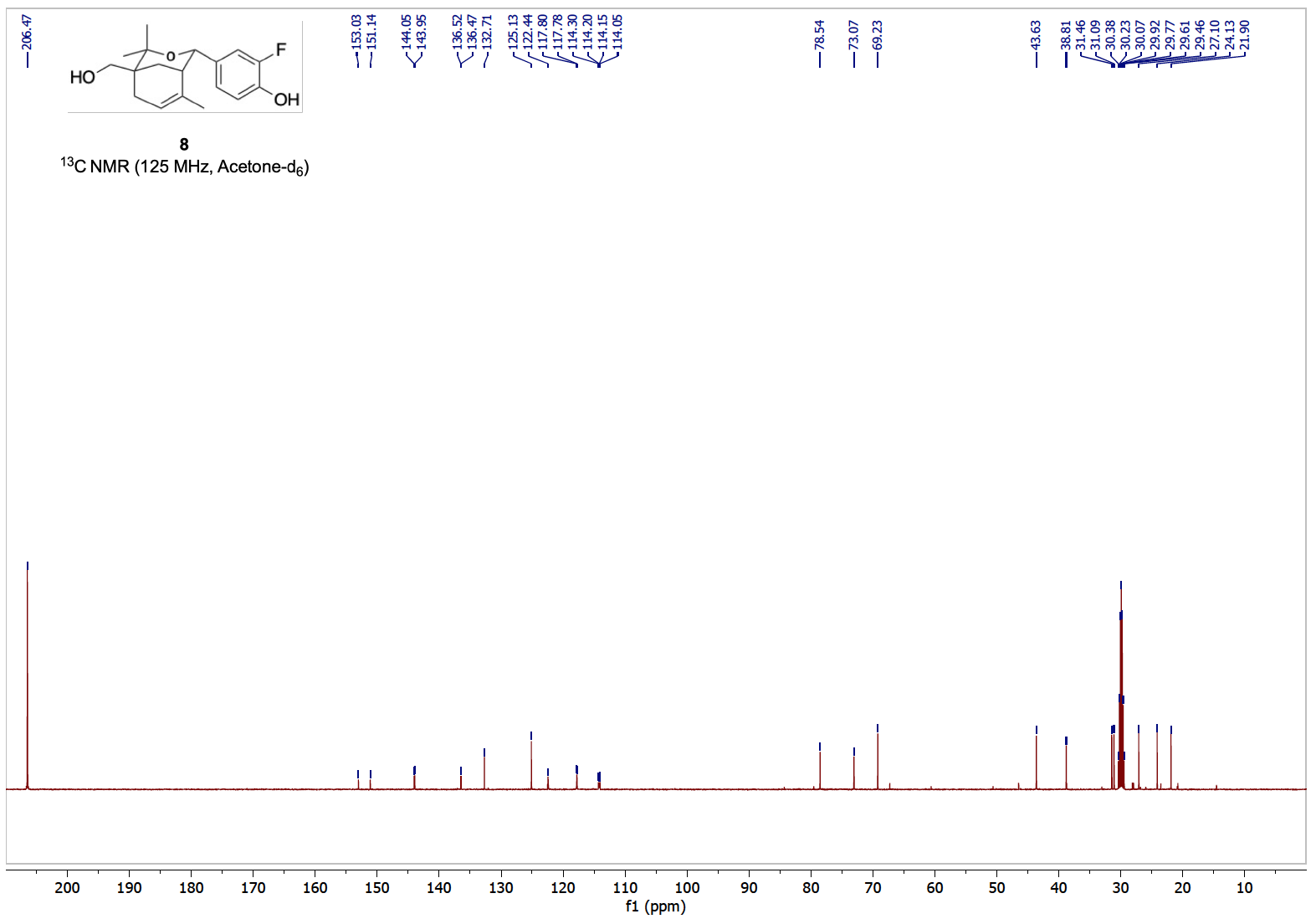

**B**

Acetone‑d_6_

(singlet)

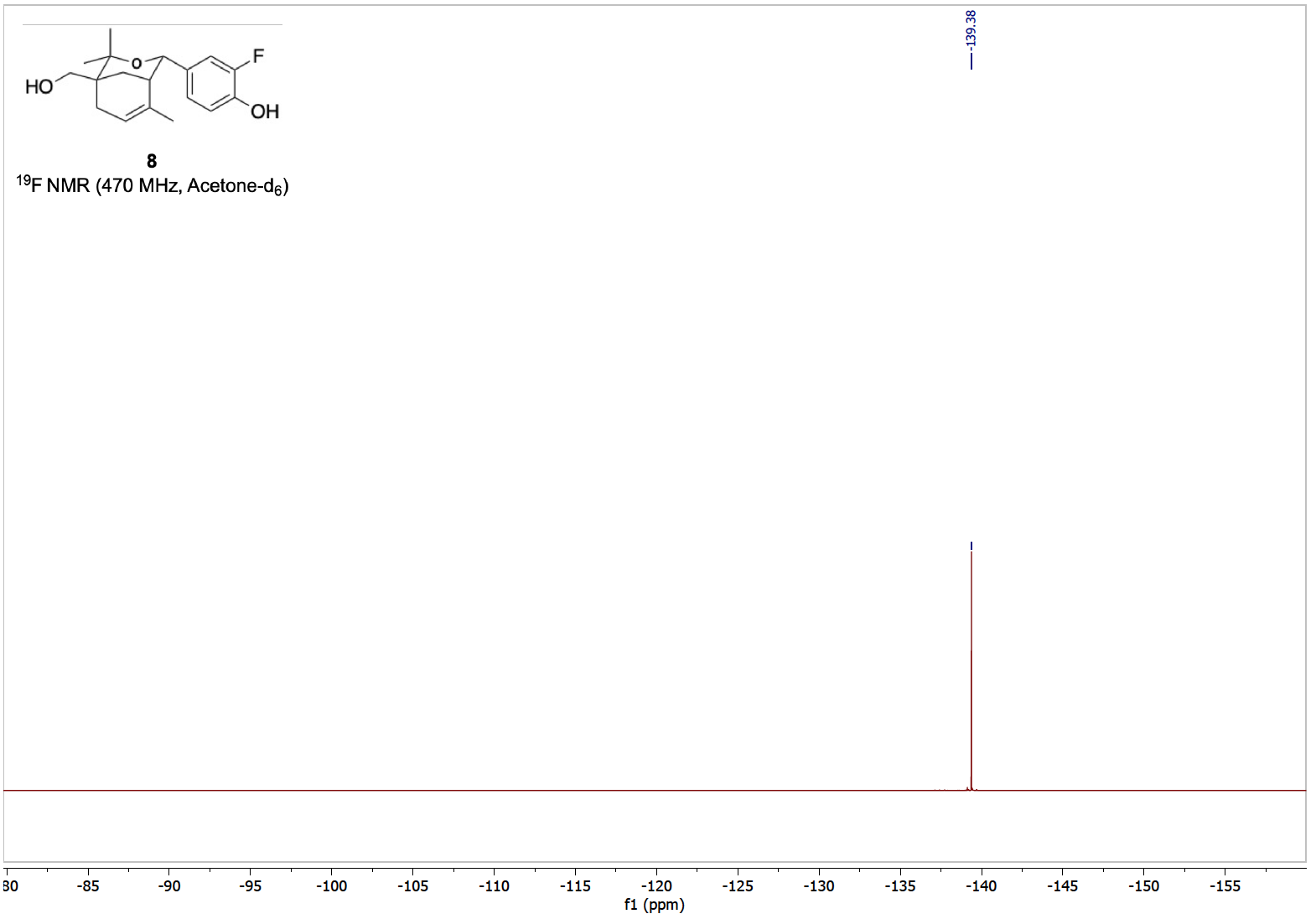

**C**

**Figure S7.** **(A)** ^1^H‑NMR (500 MHz), **(B)** ^13^C‑NMR (125 MHz) and **(C)** ^19^F‑NMR (470 MHz) of compound **8**.

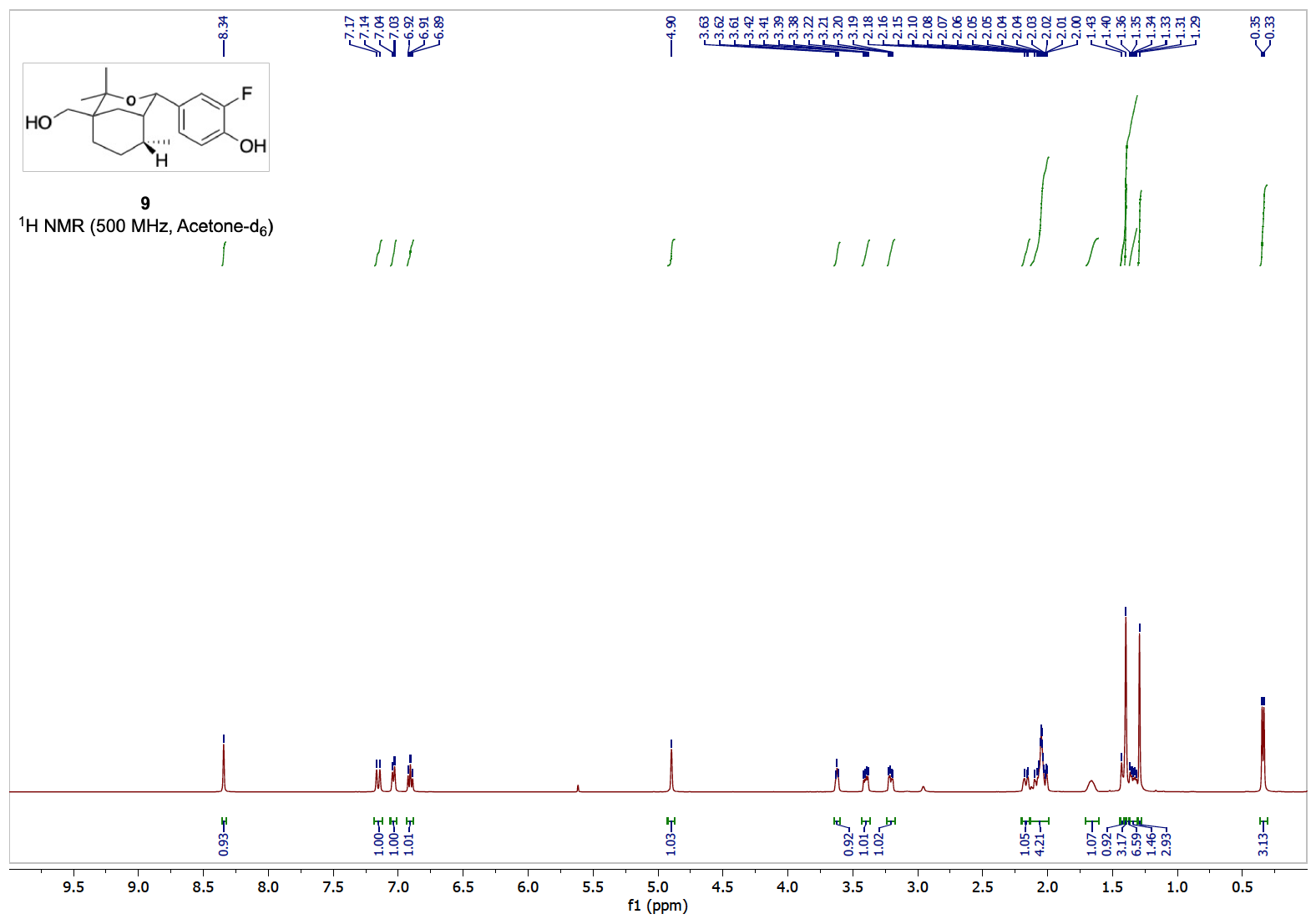

Acetone‑d_6_

(singlet)

**A**

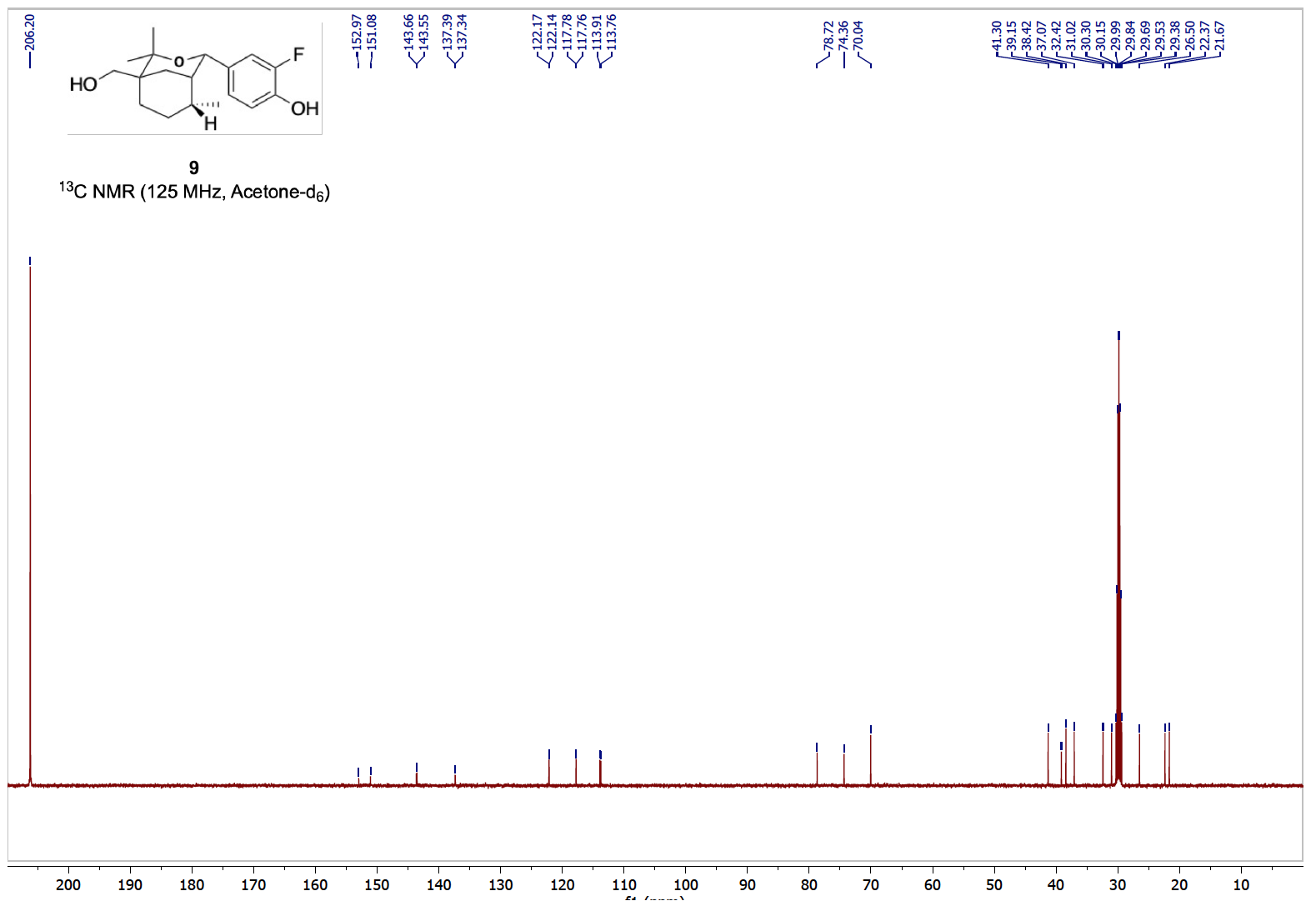

Acetone‑d_6_

(singlet)

**B**

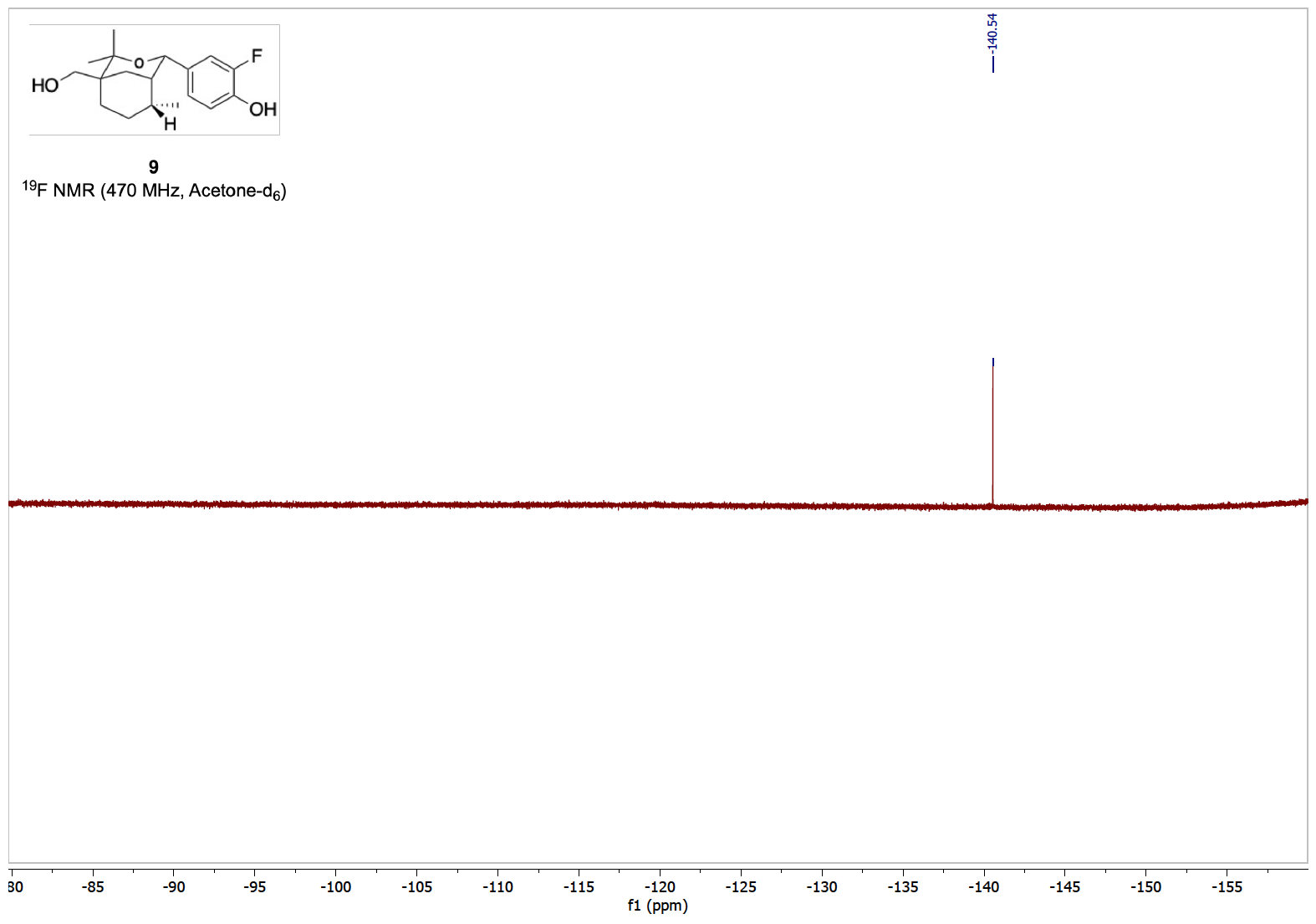

**C**

**Figure S8.** **(A)** ^1^H‑NMR (500 MHz), **(B)** ^13^C‑NMR (125 MHz) and **(C)** ^19^F‑NMR (470 MHz) of compound **9**.

**Table S1.** Summary of multi‑step syntheses of compounds **2**‑**9**.

| Compound | Transformation | Yield (%) | Notes |
| --- | --- | --- | --- |
| **2** | 11 → 2 | 67 | Prins catalyst Hf(OTf)_4_, CH_3_CN, 25 °C 22h |
| **3** | 11 → 3 | 75 | Prins catalyst Hf(OTf)_4_, CH_3_CN, 25 °C 22h |
|  | 11 → 13 | 61 | Prins catalyst Hf(OTf)_4_, CH_3_CN, 40 °C 19h |
| **4** | 13 → 4 | 90 | Hydrolysis NaHCO_3_, CH_3_OH, 25 °C 20h |
|  | *→ 14b | 85 | Diels Alder catalyst ZnCl_2_, THF, 70 °C 20h  *CH_3_COCH_2_CO_2_*^t^*Bu, CH_2_O, CH_2_C(CH_3_)CHCH_2_  Regioisomers 80:20 |
|  | 14b → 15b | 87 | Grignard CH_3_MgI, Et_2_O, 0 °C 5h  Regioisomers 82:18 |
|  | 15b → 15c | 76 | Hydrolysis CF_3_CO_2_H, DCM, 0 °C 1h |
| **5** | 15c → 5 | 52 | Prins catalyst Hf(OTf)_4_, CH_3_CN, 25 °C 18h |
|  | ** → 17 | 90 | Diels Alder: ZnCl_2_, THF, 25 °C 4h  **CH_2_C(CN)CO_2_Et, CH_2_C(CH_3_)CHCH_2_  Single Regioisomer |
|  | 17 → 18 | 83 | Grignard CH_3_MgI, Et_2_O, 0 °C 5h |
| **6** | 18 → 6 | 70 | Prins catalyst Hf(OTf)_4_, CH_3_CN, 50 °C 6h |
| **7** | 6 → 7 | 55 | Reduction LiAlH_4_, Et_2_O, 0-37 °C 48h |
|  | *** → 14a | 70 | Diels Alder catalyst ZnCl_2_, THF, 70 °C 20h  ***CH_3_COCH_2_CO_2_Et, CH_2_O, CH_2_C(CH_3_)CHCH_2_  Regioisomers 87:13 |
|  | 14a → 15a | 82 | Grignard CH_3_MgI, Et_2_O, 0 °C 5h  Regioisomers 85:15 |
|  | 15a → 16a | 83 | Prins catalyst Hf(OTf)_4_, CH_3_CN, 25 °C 24h |
| **8** | 16a → 8 | 83 | Reduction LiAlH_4_, Et_2_O, 0-40 °C 3h |
| **9** | 8 → 9 | 52 | Hydrogenation H_2_, Pd/C, EtOH 25 °C 5h |
